# Leukemic Cells Manipulate MSCs Bioelectrical Signals to Reshape the Bone Marrow Niche

**DOI:** 10.1101/2025.03.10.642319

**Authors:** Giulia Borella, Maddalena Benetton, Ambra Da Ros, Giorgia Longo, Giulia Borile, Alice Cani, Diego Lopez-Pigozzi, Mario Bortolozzi, Silvia Bresolin, Claudia Tregnago, Franco Locatelli, Martina Pigazzi

**Affiliations:** Department of Women’s and Children’s Health, Onco-Hematology lab and clinic, University of Padova, 35128 Padova, Italy; Foundation Istituto Ricerca Pediatrica Città della Speranza, 35127 Padova, Italy; Department of Physics and Astronomy "G. Galilei", University of Padova, 35121 Padova, Italy; Veneto Institute of Molecular Medicine (VIMM), 35129 Padova, Italy; Department of Pediatric Hematology and Oncology, IRCCS, Ospedale Pediatrico Bambino Gesù, Catholic University of the Sacred Heart, 00165 Rome, Italy

**Keywords:** mesenchymal stromal cells, voltage membrane potential, bioelectricity, tumor microenvironment, acute myeloid leukemia

## Abstract

Mesenchymal stromal cells (MSCs) are key components of the tumor microenvironment (TME), influencing leukemia progression through poorly understood mechanisms. We investigated the bioelectrical properties of MSCs derived from pediatric AML patients (AML-MSCs) and identified a significant depolarization of their resting membrane potential (V_mem_, −14.7mV) compared to healthy MSCs (h-MSCs, −28.5mV), accompanied by downregulation of CaV1.2 L-type calcium channel expression. AML-MSCs displayed increased spontaneous calcium oscillations, suggesting altered ion homeostasis. Notably, h-MSCs exposed to AML blasts underwent a similar V_mem_ depolarization (−11.8mV) and CaV1.2 downregulation, indicating that leukemic cells actively reprogram MSCs. Functionally, V_mem_ depolarization in h-MSCs promoted a pro-leukemic phenotype, whereas hyperpolarization of AML-MSCs restored a normal behavior. CaV1.2 over-expression by lentiviral vectors in AML-MSCs shifted V_mem_ toward hyperpolarization and partially reversed their leukemia-supportive properties, in part through CaV1.2 transfer via tunneling nanotubes.

These findings reveal that AML blasts impose a bioelectrical signature on MSCs, modulating ion channel activity to sustain a leukemic niche. Targeting this electrical reprogramming through CaV1.2 restoration represents a potential strategy to re-establish homeostasis in the bone marrow microenvironment.

## INTRODUCTION

Recent advancements in cancer research indicate that cancer is not only due to malignant cells, but involves a complex network of transformed and non-transformed cells, including nervous, endothelial, stromal, stem, and immune cells that make up the tumor microenvironment (TME)^1^. In this context of interconnected tumor cells and TME, how intra- or extra-cellular conditions and ion channels contribute to cancer is largely under-explored. Ion channels are membrane proteins that can catalyze the selective and regulated diffusion of ions across cellular membranes^2,3^. Their activity is mutually integrated into physiological processes controlling cell signaling pathways and being tightly controlled by them. In addition to well-recognized ones, like voltage-gated Na^+^/K^+^/Ca^2+^ channels in heart and neurons, novel ion channels are continuously discovered in both excitable and non-excitable cells and play important roles in developmental disorders, neurodegenerative diseases and cancer, constituting the second largest class of drug targets after G protein-coupled receptors^4,5^. Ion channels work in concert to maintain ionic homeostasis and cellular resting voltage membrane potential (V_mem_) that refers to the electric potential difference between cytoplasm and extracellular space. Their “on” or “off” state governance and ability to form electrical networks in response to inputs from local TME is under investigation^6,7^. An abnormal depolarization of V_mem_ and ion channels conductivity has been reported to play a role in cancer initiation, TME remodeling and therapy response, particularly in solid tumors^8–10^. Importantly, V_mem_ was found to impact on cellular differentiation and proliferation, with depolarization characterizing plastic, undifferentiated, proliferative cells, while increased negative V_mem_ (hyperpolarization) supporting cell differentiation and lower proliferation^7,11^. As central regulators of cellular electrical properties, ion channels might be implicated in all steps of tumorigenesis. However, in the field of ion channel discovery there are few published studies, often reporting unconfirmed data^12^. Particularly in stem and progenitor cells, including adult human mesenchymal stromal cells (MSCs), a forced depolarization has been shown to maintain cells in a stem-cell like state^13–16^, and the identification of a V_mem_ threshold to separate “normal” quiescent or resting cells from proliferative or cancerous ones is of great interest. Furthermore, observations that hyperpolarizing treatments inhibit tumor formation suggest that depolarization is a disease phenotype that is worth investigating^17^, and that the reversible nature of bioelectric tissue remodeling supports the development of innovative therapies^18–20^, with ion channels being the cancer molecular targets of approximately 15% of US FDA-approved drugs^21–23^.

Pediatric cancer is still a life-threatening disease with few new drugs in the market, particularly for acute leukemia, the most frequent tumor in childhood^24,25^. We recently attributed for the first time a role to calcium signaling and ion channels in MSCs within the leukemic bone marrow (BM) niche. Using our previously established three-dimensional (3D) BM niche model, we discovered that Lercanidipine, a voltage-gated L-type calcium channel (CaV1.2) blocker, induces functional modifications in acute myeloid leukemia (AML) derived MSCs^26^. This highlights a previously unrecognized mechanism by which ion channels activity in MSCs contributes to shaping the TME and supporting leukemia progression.

Here, we documented that non-excitable MSCs possess a resting V_mem_ that undergoes significant depolarization upon exposure to leukemic blasts. This shift is driven by altered expression of CaV1.2, a key regulator of calcium influx, which in turn promotes novel pro-leukemic functions that shape the TME.

## METHODS

### Study design

This study was designed to investigate the contribution of intracellular Ca^2+^ oscillation, observed in a 3D model of MSCs and leukemic blasts, in mediating primary MSCs V_mem_ changes (see Supplementary Table 1). Briefly, we measured Ca^2+^ flux dynamics and V_mem_ changes in h-MSCs and AML-MSCs (both derived from primary BM specimens) using the Ca^2+^ sensitive dye Fluo-4 and the voltage-sensitive dye DiBAC_4_(3) (Bis-(1,3-diethylthiobarbituricacid) trimethine Oxonol) and patch clamp. Then, we pharmacologically modulated V_mem_ and tested 32D cell line proliferation, HUVEC tube formation and lymphocytes activation, and CaV1.2 expression *in vitro*. We reverted primary AML-MSCs (namely re-AML-MSCs) by over-expressing CaV1.2 using lentiviral vectors and studied V_mem_ recovery, MSCs gene expression and functional properties, including hematopoietic stem cells (HSCs) sustainment and hematopoiesis support *in vitro* and in a transplant setting *in vivo*. Procedures involving animals were in accordance with national and international laws and policies (EEC Council Directive 86/609, OJ L 358, 12 December 1987) and “ARRIVE” guidelines (Animals in Research Reporting In Vivo Experiments), and approved by Italian Ministry of Health (512/2019-PR).

### Ethics approval and consent to participate

AML and derived MSCs are from patients in accordance with AIEOP AML 2002/01 protocol. Written informed consent to utilize biological material exceeding from the leukemia cell characterization of children enrolled in the AIEOP AML 2002/01 protocol was obtained by patients’ legal guardians and the study was performed in accordance with the Declaration of Helsinki and our local institutional review board. *In vivo* experimental procedures were performed at Venetian Oncologic Institute animal facility. Procedures involving animals were in accordance with institutional guidelines that comply with national and international laws and policies (EEC Council Directive 86/609, OJ L 358, 12 December 1987) and with “ARRIVE” guidelines (Animals in Research Reporting In Vivo Experiments). Ministry authorization approved: 512/2019-PR.

### MSCs isolation, culture and characterization

Isolation, culture, expansion and characterization of MSCs from BM were conducted as previously published^26^. Briefly, cells from BM of pediatric AML patients and healthy donors were plated at 100.000 cells/cm2 in StemMACS^TM^ MSC Expansion Media (Miltenyi Biotec, Bergisch Gladbach, DE), supplemented with 100 U/mL penicillin/streptomycin (Gibco, Life Technologies, CA, USA) and incubated at 37°C. After 1 day of culture, the non-adherent cells were removed and fresh medium was added. The resulting adherent cells have been considered to be MSCs derived from AML at diagnosis (AML-MSCS) or from healthy donors (h-MSCs). These derived MSCs were characterized as per ISCT guidelines, demonstrating positive expression for CD73, CD90, and CD105, and negative expression for CD34, CD45, and CD11b, and their multipotency to differentiate into adipogenic, osteogenic, and chondrogenic lineages. All MSCs were expanded until 90% confluence and then re-plated and expanded in larger flasks. At each passage, MSCs were trypsinized (Trypsin/EDTA Solution, Biochrom GmbH, Berlin, DE) at 37°C for 5’, harvested in medium with FBS (Thermo Fisher Scientific Waltham, MA, USA) to inactivate trypsin, spun and plated at 5 x 103 cells/cm^2^ in fresh medium. All experiments were carried out using MSCs between passages 2 to 5.

### Cell lines and primary cells cultures

AML cell lines HL-60 and SHI-1, primary AML blasts, and Human Umbilical Vein Endothelial Cells (HUVEC, Lonza, Basel, CH) were cultured as previously described^26^.

Healthy CD34^+^ cells were isolated from human cord blood (CB) mononuclear cells (PBMCs) obtained by centrifugation over Lymphoprep^TM^ (Serumwerk, Bermbrug, AG), followed by incubation with CD34 MicroBeads (Miltenyi Biotec, Bergisch Gladbach, DE). CD34^+^ cells were isolated by magnetic cell sorting and cultured in MyeloCult H5100 medium (Stem Cell Technologies, Vancouver, Canada), supplemented with cytokines (100 ng/mL, hSCF, and hFlt3L; 20 ng/mL hIL-3, hIL-6, and hG-CSF, Miltenyi Biotec).

HEK293T (DSMZ, Braunschweig, DE) cells were maintained in DMEM supplemented with 10% FBS (Thermo Fisher Scientific, Waltham, MA, USA), 2 mM glutamine (Gibco, Thermo Fisher Scientific) and 100 U/mL streptomycin/penicillin (Gibco, Thermo Fisher Scientific).

### Ca^2+^ flux imaging

Ca^2+^ flux was monitored by fluorescence imagining using the Ca^2+^ sensitive dye Fluo-4 (Invitrogen, Thermo Fisher Scientific) as previously described^26^. Time-lapse series were taken at 2-second intervals over a period of 40-400 seconds[to capture Ca^2+^ oscillations.

FIJI ImageJ software was used for image processing, including drawing of regions of interest around individual cells and calculating average pixel intensity per cell per time. Fluorescence intensity was normalized to minimum intensity values and monitored over time for each cell.

For amplitude calculation, we determined maximal (F_max_) and minimal (F_min_) fluorescence values and data are presented as the relative change in fluorescence. After monitoring baseline oscillations, Ca^2+^ levels were determined before and during the exogenous administration of KCl solution (65 mM) and then imaged for additional 50-60 min. Intracellular Ca^2+^ levels were also determined after treatment with 10 nM of Ouabain (Sigma-Aldrich-Merck, Darmstadt, DE) for 72 hours.

### V_mem_ measurements by DiBAC staining

To measure V_mem_ changes at the predetermined timepoints during cell proliferation and drug treatment, MSCs were loaded with the voltage-sensitive dye Bis-(1,3-diethylthiobarbituricacid) trimethine Oxonol (DiBAC_4_(3), Invitrogen, Thermo Fisher Scientific). The extent of DiBAC binding is inversely proportional to the magnitude of the voltage across the membrane; thus, its uptake is voltage-dependent and higher in depolarized cells. MSCs were seeded at a density of 2.5×10^3^ cells/cm^2^ in a 24-well glass-bottom plate (Ibidi GmbH, Gräfelfing, DE). After 72 hours, a fresh solution of 1 mM DiBAC_4_(3) in DMSO was prepared and diluted to 1 µM in Hank’s Buffered Salt Solution (Invitrogen, Thermo Fisher Scientific). After dye addition, cells were incubated for 20 min at 37°C, then imaged using Zeiss Observer microscope (Zeiss, Oberkochen, DE). The DiBAC_4_(3) dye was excited with a 488 nm light and fluorescence was captured at 510/530 nm. Since fluorescence intensity was quantified for each image, gain, exposure time, and offset settings of the microscope were kept constant over the duration of each experiment. Fluorescence intensity was quantified by FIJI ImageJ software.

To evaluate the influence of external Ca^2+^ mobilization on V_mem_ modulation, MSCs were treated with the extracellular calcium chelator EGTA (2 mM, Sigma-Aldrich-Merck), whereas the effect of intracellular calcium was studied by using the calcium chelator BAPTA-AM (50 µM, Selleckchem, Houston, TX, USA).

### Intracellular V_mem_ recordings through perforated patch clamp

V_mem_ was recorded in AML-MSCs and h-MSCs cultured either alone or with AML blasts after 24-72 hours from seeding. Cells were resuspended over a glass previously coated with fibronectin 40 µg/mL (Corning Life Sciences, NY, USA) to a chamber at room temperature (RT). The extracellular solution contained 150 mM NaCl, 5 mM KCl, 2 mM CaCl_2_, 1 mM MgCl_2_, 2 mM sodium pyruvate, 10 mM HEPES, 5 mM glucose with 310 Osm and pH adjusted to 7.4 using NaOH. DiBAC 1 µM was added while waiting 15 min for cells to precipitate in the coverslip. We used borosilicate glass capillary tubing (1.5 mm OD, Warner Instrument Corp., Hamden, CT) pulled with a vertical puller (Model P-830, Narishige International, Japan) resulting in pipette resistances in the range of 6-8 MΩ. Pipettes were filled with an intracellular solution containing 115 mM potassium aspartate, 10 mM NaCl, 10 mM KCl, 1 mM MgCl_2_, 10 mM HEPES, 4 mM BAPTA with 285 Osm and pH adjusted to 7.2 with KOH. Gramicidin A 0.1 mg/mL (Sigma-Aldrich-Merck) was added freshly each day of recordings.

Cell currents were amplified by a Multiclamp 700b amplifier (Molecular Devices, CA, USA) and recorded by the pClamp software (version 10.4, Molecular Device) after digitalization at 10 KHz by a Digidata 1550A (Molecular Devices). Data analysis was performed by custom-made software in MatLab 2019b (MathWorks Inc., MA, USA). DiBAC fluorescence was excited by a 470-nm collimated LED (Thorlabs, NJ, USA) and CD45-A568 fluorescence with a 565-nm collimated LED (Thorlabs) filtered with 535/50 BP and 610/LP filters, respectively. The fluorescence light was collected using an upright wide-field microscope (BX51WI, Olympus, Tokyo, Japan) equipped with an infinity-corrected water immersion objective (40x, 0.8 NA, Olympus) and a sCMOS cooled camera (PCO.edge 5.5) at 0.1 Hz frame rate.

To achieve perforation, the patched cell was held at 0 mV in cell attached configuration until Gramicidin A allowed to record intracellularly, normally after 10 to 15 min. DiBAC initial fluorescence was then measured (F_0_) before switching to current clamp at 0 pA in order to measure the resting V_mem_. In this configuration, DiBAC fluorescence (F) was measured once stabilized for at least 5 min, and then holding again the cell at 0 mV to check for F_0_ reliability. DiBAC relative fluorescence variation, ΔF/F_0_=(F-F_0_)/F_0_, was found to be proportional to V_mem_ in first approximation.

### Pharmacological V_mem_ modulation

Several drugs were employed to modulate V_mem_ in MSCs: (1) Na^+^/K^+^-ATPase-inhibitor Ouabain (5-25 nM, Sigma-Aldrich-Merck) was added to the medium from a fresh 10 mM stock solution in DMSO; (2) the concentration of extracellular K^+^ was increased by adding potassium gluconate (K^+^ gluconate, Sigma-Aldrich-Merck) to the medium to final concentrations of 10–80 mM; (3) the chloride channel (ClC-2) activators Lubiprostone (Lubi, Selleckchem) and Ivermectin (IVM, Sigma-Aldrich-Merck) were added to the medium to final concentrations of 0.1-1 µM and 1-50 µM, respectively, from 10 mM stock solutions in DMSO. For all experiments, MSCs were seeded at 5×10^3^ cells/cm^2^ in StemMACS^TM^ MSC Expansion Media (Miltenyi Biotec); the day after, cells were treated with the depolarizing/hyperpolarizing agent for 72 hours. In order to evaluate the reversibility of the effect, drug washout procedure was performed and after 72 hours its effects were evaluated.

In order to explore the influence of V_mem_ in CaV1.2 modulation, MSCs pre-treated with depolarizing/hyperpolarizing agents for 72 hours were then exposed to the CaV1.2 blocker lercanidipine (10 µM, Sigma-Aldrich Merck) for 48 hours.

### Cell proliferation assay

MSCs proliferation was evaluated using PrestoBlue^TM^ Cell Viability Reagent (Invitrogen, Thermo Fisher Scientific) following the manufacturer’s guidelines. Briefly, 2,5×10^3^ MSCs were cultured in 100 μL of proper medium in sterile 96-well plates and incubated at 37°C, then at the indicated timepoints 10 μL of PrestoBlue reagent were added. Plates were incubated for 1 hour at 37°C and absorbance was recorded using a Spark^®^ spectrophotometer (TECAN, Männedorf, CH) at 570 nm with a reference wavelength of 600 nm.

### MSCs-32D co-culture and differentiation evaluation

IL-3-dependent 32D murine hematopoietic precursors cells (DSMZ) were cultured for 72 hours on a layer of depolarized or hyperpolarized MSCs, as previously described^26^.

To assess 32D cell differentiation, 32D cells in suspension were collected and centrifugated to allow attachment to the slice, then stained by May–Grünwald Giemsa method.

### HUVEC Tube Formation Assay with conditioned medium of MSCs

HUVEC Tube Formation Assay was conducted as published previously^26^, by using conditioned media derived from depolarized and hyperpolarized-MSCs.

### Lymphocytes activation assay

Anti-inflammatory potential of depolarized- and hyperpolarized-MSCs was tested evaluating the activation of proliferating lymphocytes, as we previously described^26^.

### Human IL-6 ELISA Assay

IL-6 protein levels in culture supernatants were measured using ELISA (Thermo Fisher Scientific), following manufacturer’s instructions. Absorbance A450 values were measured using a microplate reader (Spark^®^, TECAN).

### RNA isolation and quantitative real-time PCR

Total RNA was isolated using Trizol (Invitrogen, Thermo Fisher Scientific). One μg of RNA was reverse-transcribed into cDNA using the SuperScript II system (Invitrogen, Thermo Fisher Scientific) according to the manufacturer’s instructions. Expression of mRNA was measured by Real Time PCR (RQ-PCR) on an ABI 7900HD platform (Applied Biosystems, Thermo Fisher Scientific) using the SYBR Green PCR master mix (Applied Biosystems, Thermo Fisher Scientific) and normalized on *GUSB* housekeeping gene, using the 2^-ΔΔCt^ method. Primers sequences for the selected genes are reported in the following table:

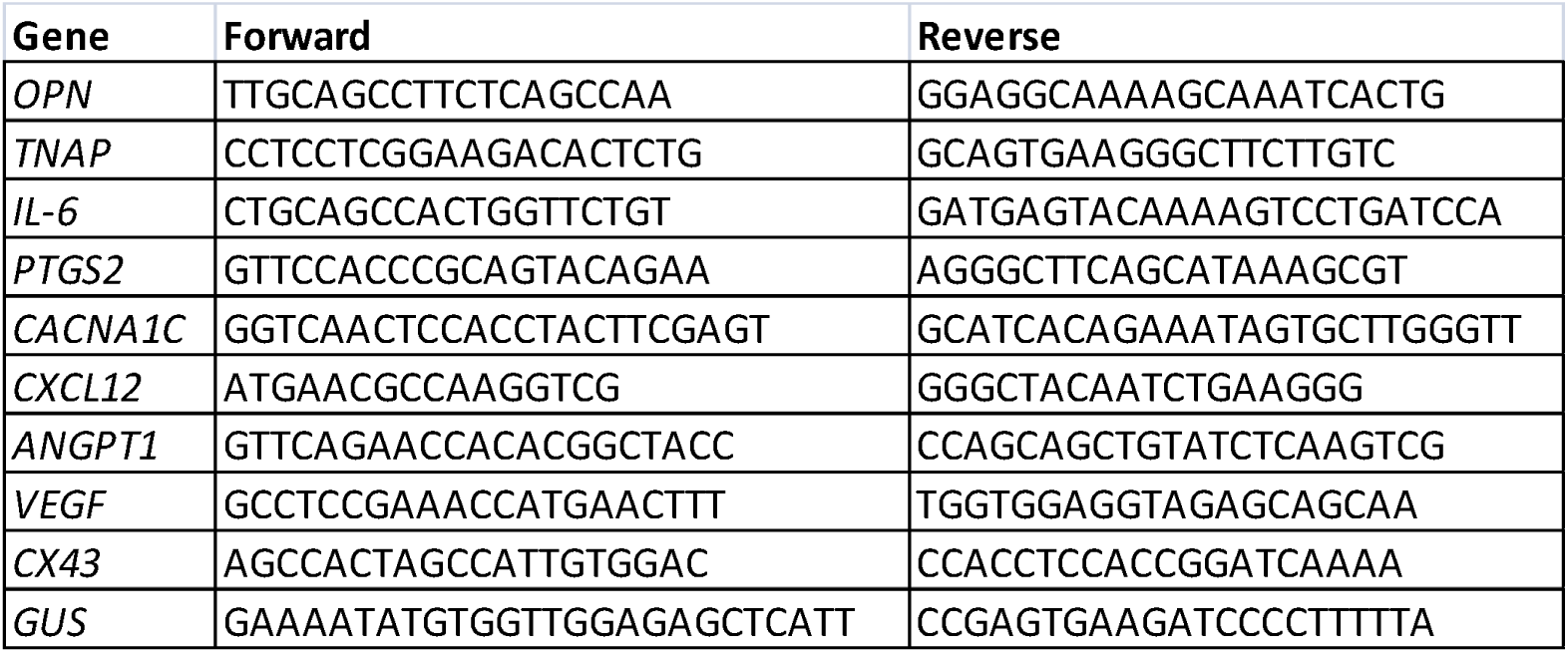

### AML-induced MSCs co-culture model

h-MSCs were plated at 5×10^3^/cm^2^ and, after 24 hours, co-cultured with AML primary blasts or AML cell lines (ratio 1:10) for a timing between 30 min and 7 days in AML cells proper medium. After being co-cultured with AML cells, h-MSCs have been named “induced AML-MSCs” (iAML-MSCs). As control, h-MSCs were co-cultured with CD34^+^ cells from CB.

The role of cell-cell contact was investigated by silencing Connexin-43 (Cx-43, Ambion, Thermo Fisher Scientific) in AML blasts and h-MSCs or by treatment with Carbenoxolone (CBX, 100 µM, Sigma-Aldrich-Merck) for 4 days of co-culture. The role of AML blasts secretome was evaluated by using a 0.4 µm polycarbonate Transwell inserts (Corning Life Sciences) to separate AML blasts to MSCs for 4 days.

### Primary cell transfection

Cx-43 was silenced in AML primary blasts and h-MSCs, using electroporation or calcium phosphate methods as previously described ^26^.

### Intracellular staining for flow cytometry analysis

Cells were fixed in 4% formaldehyde for 10 min, then permeabilized in 0.1% Tween20 (Sigma-Aldrich-Merck) for 20 min, blocked with 3% bovine serum albumin (BSA, Sigma-Aldrich-Merck) for 30 min, and incubated with 1:500 primary anti-human CaV1.2 antibody (Alomone labs, Jerusalem, Israel) in 3% BSA+20% FcR Blocking Reagent (Miltenyi Biotec) for 30 min. Next, cells were incubated with Alexa Fluor 488 Goat Anti-Rabbit IgG (H+L) secondary antibody (1:500, Thermo Fisher Scientific) for 30 min at RT, washed and analyzed with CytoFLEX (Beckman Coulter).

### Transient plasmid transfection of HEK293T

Cells were grown as a monolayer to confluence up to 70% in a 35 mm Petri dish prior to transfection. Transient transfection was performed using calcium phosphate method, using 0.33 μg of plasmid encoding CaV1.2-mCherry (*pLV[Exp]-mCherry-hPGK>hCACNA1C*, Vector Builder Inc., IL, USA).

### MSCs transduction with lentiviral vectors

MSCs at passage 1 were seeded at a density of 1.5×10^5^ cells/cm^2^ in 6-well plates in a final volume of 2 mL. The day after, the vial containing the LV (Vector Builder Inc., Chicago, IL, USA) was thawed and added to MSCs with protamine sulfate (100 µg/mL in DMEM, Sigma-Aldrich-Merck), at a multiplicity of infection (MOI) of 50. The volume of LV used was calculated as follows: MOI x N/TU, where TU is the amount of functional LV particles per unit of volume, and N is the number of cells. The plate was incubated at 37°C for 18 hours and then cells were extensively washed.

MSCs were transduced in parallel with CaV1.2-mCherry (*pLV[Exp]-mCherry-hPGK>hCACNA1C*) and control GFP-mCherry (*pLV[Exp]-EGFP:T2A:Puro-EF1A>mCherry*, Vector Builder) LVs. The efficiency of transduction was measured after 7 and 14 days by detecting the mCherry positivity by flow cytometry on the BD FACSCelesta^TM^ (BD). CaV1.2 over-expression was verified by RQ-PCR and by flow cytometry after 14 days from transduction. Un-transduced (UT) MSCs, incubated only with protamine sulfate, were used as control.

### Immunofluorescence

Cells were fixed in 4% formaldehyde for 10 min, washed in PBS 1X, permeabilized with 0.5% Triton X-100 (Sigma-Aldrich-Merck) for 20 min, washed in PBS 1X, blocked with 1% BSA. After 30 min, cells were subjected to immunofluorescence using the anti-CaV1.2 antibody (1:200, Alomone labs) overnight at 4°C. Primary antibody incubation was followed by incubation with appropriated fluorochrome-conjugated secondary antibodies 2 hours at RT. Cells were counterstained with 4[,6-diamidino-2-phenylindole (DAPI; 1:10000; Sigma-Aldrich-Merck) to label nuclei and with Alexa Fluor™ 647 Phalloidin (Invitrogen, Thermo Fisher Scientific) to label F-actin. Images were acquired using Zeiss Observer microscope (Zeiss). Fluorescent intensity of MSCs was quantified by FIJI ImageJ software. Data were represented as mean pixel intensity ± SEM.

### Gene expression analysis of MSCs

We analyzed RNA collected from 6 h-MSCs, 21 AML-MSCs, 2 AML-MSCs UT, and 7 Re-AML-MSCs after 3 passages of *in vitro*. RNA quality was assessed on an Agilent2100 Bioanalyzer (Agilent Technologies, CA, USA). Then, 100 ng of total RNA were labeled and hybridized to GeneChip™ Human Genome U133 Plus 2.0 Array (Applied Biosystems, Thermo Fisher Scientific) for 16 hours at 45°C using a rotational oven and washed according to Affymetrix standard protocols using a GC450 Fluidics Station (Applied Biosystems, Thermo Fisher Scientific). The GeneChip were scanned with an Affymetrix 7G scanner and CEL files generated were normalized by robust multi-array averaging (RMA) algorithm with Affy-R package (www.r-project.com). ComBat function in SVA package was used to eliminate batch effect among experiments. Principal Component Analysis (PCA) using probe sets with more than 90% of variance variability was obtained.

The datasets generated and analyzed during the current study are available in the GEO repository, with the access code GSE248681.

### 3D culture system setup

The 3D bone marrow niche model consists of a biomimetic scaffold composed of hydroxyapatite and collagen type I. Scaffolds seeding with primary MSCs and AML cells was conducted as previously reported^26^.

### V_mem_ analysis in 3D

To measure V_mem_ in 3D after one week of culture, the voltage-sensitive dye DiBAC_4_(3) was diluted to 1 µM and added to the culture media for 20 min at 37°C. To stain for nuclei, Hoechst 33342 (Thermo Fisher Scientific) at 8 µM was subsequently added for 10 min at 37°C. After loading, scaffolds were immediately imaged using Zeiss Observer microscope (Zeiss, Oberkochen, DE). Fluorescence intensity was quantified by FIJI ImageJ software.

### 3D-AML culture staining

The 3D culture was fixed with 4% Paraformaldehyde for 30’ at 4°C, then cells were permeabilized *in situ* with 0.5% TritonX-100 and 5% bovine serum albumin (BSA) (both from Sigma-Aldrich-Merck) in PBS 1x for 1 hour at room temperature (RT). For immunofluorescence staining, the primary antibody (Ki-67 1:200, Dako - Agilent Technologies, Santa Clara, CA, USA) was diluted in 0.1% TritonX-100 and 1% BSA in PBS 1x and incubated for 24 hours at room temperature in agitation. For immunofluorescence detection, the samples were incubated with Alexa488-conjugated secondary antibody (Thermo Fisher Scientific, 1:200 in PBS 1x with 1% BSA) overnight at room temperature. F-actin and nuclei were labeled with phalloidin-Alexa647 (1:500, Thermo Fisher Scientific) and 4’,6-diamidino-2-phelylidine dihychloride (DAPI SigmaAldrich-Merck, 1:1000) respectively, in PBS 1x, for 3 hours at room temperature. After 3 washes the samples were transferred on a bottom glass chamber slides (IBIDI, Gräfelfing, DE) for confocal imaging using the Zeiss 800 Confocal microscope (Zeiss, Oberkochen, DE).

### MSCs and healthy CD34^+^ co-culture

MSCs and CD34^+^ isolated from CB of healthy donors were co-cultured as previously described^26^. 1×10^5^ CD34^+^ were plated in 500 µL of MSCs conditioned medium on a MSCs layer. After 5 days, CD34^+^ were collected and counted using Trypan Blue. The percentage of high proliferative index was evaluated by flow cytometry analysis, after incubating with Ki-67-488 (Invitrogen, Thermo Fisher Scientific) for 10 min at RT in the dark, using CytoFLEX (Beckman Coulter). CD34^+^ cells alone (stroma-free, SF) or on a layer of h-MSCs in HSCs proper medium were used as controls.

### Transplantation of human CB CD34^+^ cells and MSCs in mice

Adult NSG (7 weeks old) or NOG-EXL female mice were sub-lethally irradiated (1.5 Gy) and transplanted via tail vein injection with 1×10^5^ human CD34^+^ cells alone or in combination with 1×10^6^ human MSCs. Peripheral blood (PB) samples were collected weekly after 3 weeks from human cell co-infusion. The presence of human cells was determined by flow cytometry using the following antibodies: mouse CD45 PE-Cy7 (Thermo Fisher Scientific), human CD45 ECD (Beckman Coulter). All samples were run on a CytoFLEX (Beckman Coulter) and at least 10.000 events were recorded.

### CFSE staining in CD34+ cells

For cell proliferation assay, 1×10^6^ CD34+ cells were resuspended in 1 mL of RPMI 1640 (no supplements added) with the addition of CellTrace™ Violet Cell Proliferation Kit at the final concentration of 5 μM (Life Technologies, Thermo Fisher Scientific). After a 20-minute incubation at 37°C, 5 mL of RPMI 1640 (supplemented with 10% FBS, 2 mM glutamine and 100 U/ml streptomycin/penicillin) were added and cells were incubated for 5 additional minutes at 37°C, then washed and injected in mice.

### *In vivo* tracking of MSCs and CD34+ cells

Adult NOG-EXL (NOD.Cg-*Prkdc^scid^ Il2rg^tm1Sug^* Tg(SV40/HTLV-IL3,CSF2)10-7Jic/JicTac (7 weeks old) female mice were sub-lethally irradiated (1.5 Gy) and transplanted via tail vein injection with 1×10^5^ human CD34^+^ cells CFSE-stained with 1×10^6^ human MSCs transduced to express the mCherry reporter gene for MSCs tracking. Mice were sacrificed at 24, 48 and 72 hours post injection to evaluate the presence of MSCs in the different mice organs. At 72 hours, CD34+ cell proliferation was evaluated in murine bone marrow. Cells were monitored for fluorescence by flow cytometry on the BD FACSCelesta^TM^ (BD) cytometer (Becton Dickinson). Data were analyzed using FlowJo v10 software (FlowJo LLC, Becton Dickinson).

### Statistics

Statistical analyses were performed using Prism6 (Graph Pad Software Inc., La Jolla, CA, USA). All experiments were performed at least in duplicate, and results were presented as mean ± SEM. Statistical significance of differences between two groups was evaluated by applying the Student’s t-test, after a preliminary testing of normal data distribution. When comparing more than two groups, we applied the Holm-Šídák correction for multiple statistical hypotheses testing. Results were considered statistically significant when *p*-value was <0.05 (*), <0.01 (**), <0.001 (***), and <0.0001 (****).

## RESULTS

### Ca^2+^ flux dynamics are altered in depolarized AML-MSCs

We previously characterized MSCs derived from pediatric AML samples collected at diagnosis (AML-MSCs) highlighting pro-oncogenic features and reduced CaV1.2 expression with respect to MSCs derived from age-matched healthy donors (h-MSCs), making them sensitive to the Ca^2+^-channel blocker Lercanidipine^26^. This observation suggested that Ca^2+^ homeostasis could play a critical role in modifying MSCs behavior within the BM niche. Using Fluo4-AM in h-MSCs and AML-MSCs, we observed that AML-MSCs increased spontaneous Ca^2+^ oscillations (*p*<0.05, Fig. 1A), which had a higher amplitude than in h-MSCs (*p*<0.05, Fig. 1B). In addition, in response to high-KCl depolarization stimulus, AML-MSCs significantly reduced their extracellular Ca^2+^ uptake with respect to h-MSCs (*p*<0.0001, Fig. 1C). Since several works suggested that changes in intracellular Ca^2+^ handling are associated with CaV1.2 and V ^9,27^, we investigated MSCs V_mem_ by loading AML- and h-MSCs with the voltage-sensitive dye DiBAC, whose uptake is higher in depolarized cells, in response to KCl depolarization stimulus. Results showed that AML-MSCs showed a different DiBAC fluorescence intensity uncovering a different resting V_mem_ (Fig. 1D) and, upon depolarizing KCl stimulus, DiBAC fluorescence increased in a time-dependent manner. Thus, we measured V_mem_ of several h-MSCs (n=7) and AML-MSCs (n=14) confirming AML-MSCs depolarization compared to h-MSCs (*p*<0.0001, Fig. 1E, Supplementary Fig. 1A). We documented that h-MSCs V_mem_ ranged from 15,500 to 26,000 (DiBAC mean pixel intensity), whereas AML-MSCs from 26,000 to 38,500, at day 3 of culture (Supplementary Table 2). We validated V_mem_ measurements by patch clamp, confirming AML-MSCs depolarization with respect to h-MSCs (V_mem_ average= −14.7 mV ± 3.36 mV and −28.5 mV ± 2.58 mV respectively, *p*<0.01, Fig. 1F-G, Supplementary Fig. 1B). These latter findings showed, to the best of our knowledge, for the first time, that non-excitable MSCs acquire a different V_mem_ at leukemia onset, rendering them AML-MSCs.

**Fig. 1.**
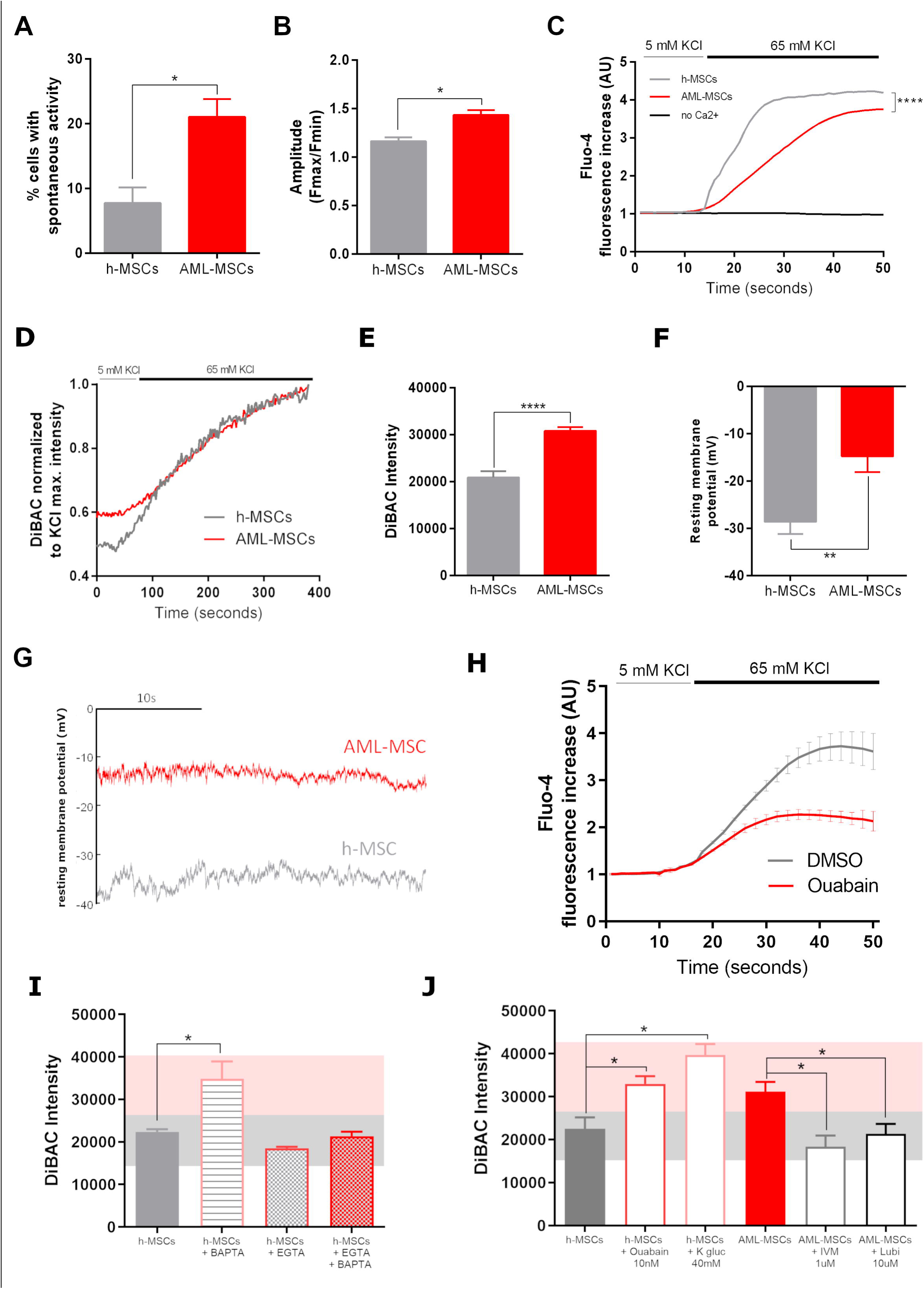
Ca^2+^ flux and resting V_mem_ are altered in AML-MSCs. (A) Percentage of spontaneous Ca^2+^ oscillating h-MSCs and AML-MSCs cells loaded with calcium indicator Fluo-4 AM probe (n=3, t-test). (B) Average amplitude of calcium oscillations in h-MSCs and AML-MSCs loaded with Fluo-4 AM probe (n=3, 7 and 50 cells per sample respectively, t-test). (C) Intracellular calcium increases in h-MSCs and AML-MSCs, loaded with Fluo-4 AM probe and stimulated with KCl (65 mM), in the presence/absence of external Ca^2+^ (n=3, t-test at maximum h-MSCs fluorescence increase. (D) DiBAC fluorescence intensity in h-MSCs and AML-MSCs loaded with the voltage-sensitive dye DiBAC and stimulated with KCl (65 mM). Fluorescence intensity data were normalized to KCl maximum intensity value (n=4, t-test at 200, 300 and 400 seconds, *p*=not significant). (E) Fluorescence intensity calculated from DiBAC fluorescence staining in AML-MSCs (n=14) and h-MSCs (n=7) (25 cells/image, 4 images, t-test). (F) Histogram of average resting membrane potentials (mV) measured by patch clamp in AML-MSCs (n=5, 11 cells) and h-MSCs (n=4, 10 cells, t-test). (G) Representative traces of two cells where resting membrane potential was recorded in current clamp configuration. (H) Intracellular calcium increases in h-MSCs treated or not with the depolarizing agent Ouabain 10 nM, and then loaded with Fluo-4 AM probe and stimulated with KCl (n=4, 100 cells, AU: arbitrary unit, t-test performed at every other second, being significant from 30 to 50 seconds). (I) DiBAC fluorescence intensity in h-MSCs treated with BAPTA (50 µM) or EGTA (2 mM) alone or in combination (n=3, t-test comparing all treated groups *versus* h-MSCs). (J) DiBAC fluorescence intensity in h-MSCs treated with Ouabain (10 nM, n=4) or K^+^ gluconate (40 mM, n=3, t-test comparing treated groups *versus* h-MSCs) and in AML-MSCs treated with Ivermectin (IVM, 1 µM, n=3) or Lubiprostone (Lubi, 10 µM, n=5, t-test comparing treated groups *versus* AML-MSCs) for 72 hours. All histograms show mean ± SEM; \**p*-value <0.05, \*\**p*-value <0.01, \*\*\*\**p*-value <0.0001.

To determine the contribution of Ca^2+^ dynamics in MSCs depolarization, we treated h-MSCs with Ouabain, a Na^+^/K^+^-ATPase inhibitor that induce a tonic depolarization, observing that Ca^2+^ uptake was significantly lowered compared to untreated cells (Fig. 1H), confirming an association between membrane depolarization and Ca^2+^ levels. In addition, we treated h-MSCs with the Ca^2+^ selective intracellular and extracellular chelators, BAPTA-AM and EGTA respectively, and monitored V_mem_ changes showing that treatment with BAPTA shifted DiBAC intensity towards positive voltages-depolarization (*p*<0.05, Fig. 1I), whereas in presence of EGTA, which blocks Ca^2+^ influx, V_mem_ remained constant. These results demonstrate V_mem_ dependence on Ca^2+^ influx dynamics and supports a bi-directional interplay between Ca^2+^ handling, the central coordinator of excitable cells functions, and MSCs depolarization.

We modulated V_mem_ by ion channel blockers or activators to promote bioelectric changes without affecting cell viability, and simultaneously tracked the corresponding voltage changes. H-MSCs were treated with the depolarizing agent Ouabain at 5, 10, 25 nM, or with K^+^ gluconate, a blocker of K^+^ channels, at 10, 40, 80 mM, for 3 days. Both treatments induced depolarization with increasing concentration (Supplementary Fig. 1C-D). In particular, the depolarizing treatment with 10 nM Ouabain or alternatively 40 mM K^+^ gluconate displayed a time-dependent effect, reaching a V_mem_ similar to that of AML-MSCs (pink bar headband, *p*<0.05, Fig.1J) and after 3 days of drug washout V_mem_ maintained depolarized status (Supplementary Fig. 1E).

On the contrary, we modulated AML-MSCs V_mem_ by using hyperpolarizing agents, Ivermectin (IVM), an activator of glutamate-gated chloride channels, at 0.1 and 1 µM, and Lubiprostone (Lubi), an activator of ClC-2 chloride channels, at 1, 10, 50 µM, for 3 days. Interestingly, we found that DiBAC fluorescence significantly decreased with 10 µM of Lubi or with 1 µM of IVM from 24 hours of treatment, and persisted after 72 hours and after the washout, indicating AML-MSCs V_mem_ hyperpolarization by drug exposure (grey headband, *p*<0.05, Supplementary Fig. 1F-G-H).

### MSCs V_mem_ pharmacological depolarization induced functional reprogramming

Considering these findings, we hypothesized that depolarization of MSCs could be the “first hit” to transform MSCs at leukemia onset, inducing their reprogramming as we previously documented^26^. To address this hypothesis, we investigated MSCs functions *in vitro* after exposure to depolarizing agents. In detail, we observed that depolarizing treatments induced a higher h-MSCs proliferation (Ouabain, 2.5-fold; K^+^ gluconate, 1.9-fold, *p*<0.05, Fig. 2A). Furthermore, depolarized MSCs supported the cytokine-dependent 32D cell growth without IL-3 (Fig. 2B-C) at the same extent of AML-MSCs, suggesting that depolarization induced novel MSCs capabilities. Moreover, we documented 32D cell differentiation after co-culture with AML-MSCs or h-MSCs treated with depolarizing agents, shown by a reduced nuclear size, enlarged cytoplasm, and a loss of dark-violet staining, when compared to h-MSCs or with standard IL-3 (Supplementary Fig. 2). Then, considering that we previously documented an impaired AML-MSCs anti-inflammatory potential compared to h-MSCs^26^, we confirmed a reduction of h-MSCs immunomodulation activity after the exposure to depolarizing agents, as observed in the positive control AML-MSCs (*p*<0.05, Fig. 2D). Accordingly, PHA-activated CD3^+^ T cells cultured on a layer of h-MSCs reduced their CD69 and CD25 expression, whereas the depolarized h-MSCs and the AML-MSCs did not induce down-regulation of these T-cell activation markers, supporting the hypothesis that depolarization reduced MSCs immunomodulatory properties (Fig. 2E). Additionally, we documented *IL-6* gene over-expression in the depolarized h-MSCs (*p*<0.01, Fig. 2F), together with a significantly higher release of IL-6 in h-MSCs treated with K^+^ gluconate (1.7-fold, *p*<0.05, Fig. 2G). In addition, we monitored h-MSCs expression of key osteo-progenitor genes after depolarization treatment, finding a significant increase of *TNAP* and *OPN* expression levels as observed in AML-MSCs (Fig. 2H-I). In depolarized h-MSCs we observed a significant up-regulation of the inflammation-related *PTGS2* gene (Fig. 2J), previously identified as the top differentially over-expressed gene between h-MSCs and AML-MSCs^26^. Altogether, V_mem_ depolarization drives a reprogrammed stroma toward the enhancement of an inflamed leukemic niche.

**Fig. 2.**
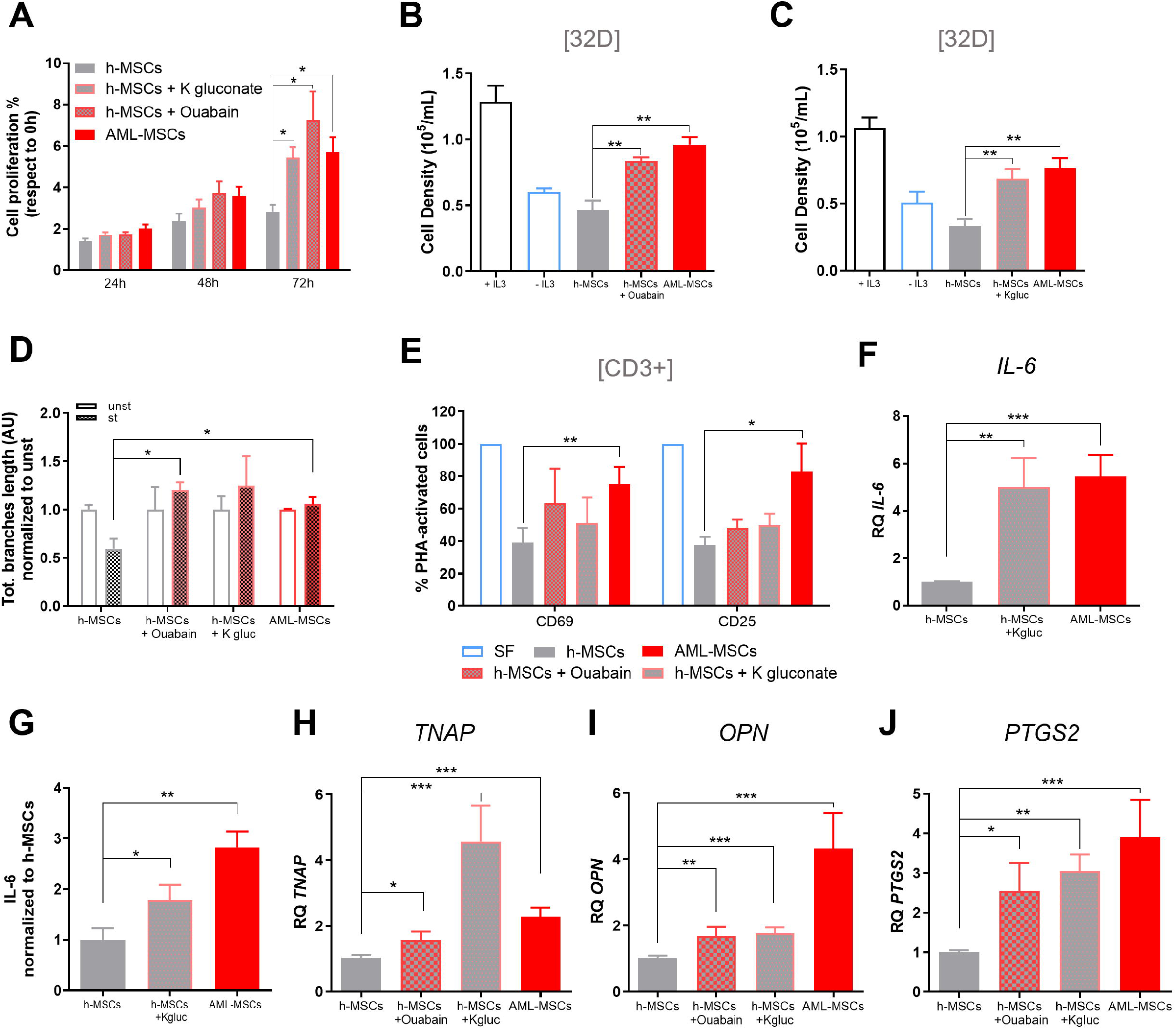
V_mem_ pharmacological depolarization induce an AML-MSCs-like phenotype in h-MSCs. (A) Cell proliferation of AML-MSCs, h-MSCs, and h-MSCs treated with 40 mM K^+^ gluconate or 10 nM Ouabain by Presto Blue assay (n=5). Data were normalized to time 0-hour samples (t-test comparing all groups *versus* h-MSCs). (B-C) Cell density of murine IL-3–dependent 32D cell line cultured in the presence (black bar) or absence of IL-3 (blue bar), compared with 32D cell line cultured on a layer of AML-MSCs (red bar), h-MSCs (grey) or h-MSCs pre-treated for 72 hours with Ouabain (B, n=5) or with K^+^ gluconate (C, n=5). T-test was performed to compare treated groups or AML-MSCs *versus* h-MSCs, with ±IL3 used as experimental control. (D) Total branches length of HUVEC tubes by using conditioned medium derived from AML-MSCs (n=7) and h-MSCs pre-treated or not for 72 hours with Ouabain or K^+^ gluconate and then stimulated (st) or not (unst) with a pro-inflammatory cytokine cocktail (hIL-1β, hIL-6, and hTNF-α) for 24 hours; tube formation was evaluated after 4 hours and normalized to unst condition (AU: arbitrary unit, n=4, t-test comparing treated or AML-MSCs stimulated groups *versus* stimulated h-MSCs). (E) Percentage of PHA-stimulated CD3^+^ T cells expressing CD69 and CD25 after 72 hours of co-culture with AML-MSCs and h-MSCs pre-treated for 72 hours with Ouabain or K^+^ gluconate, relative to SF condition (without MSCs) (n=4, t-test comparing treated or AML-MSCs groups *versus* h-MSCs). (F) Relative expression measured by RQ-PCR of *IL-6* (interleukin-6) in h-MSCs pre-treated or not for 72 hours with K^+^ gluconate (n=4) and in AML-MSCs (n=3, t-test comparing treated or AML-MSCs groups *versus* h-MSCs). (G) IL[6 protein secretion levels (pg/mL), measured by ELISA, in AML-MSCs (n=24) or in h-MSCs pre-treated or not for 72 hours with K^+^ gluconate (n=4 or n=6 respectively), relative to h-MSCs untreated condition (t-test comparing treated or AML-MSCs groups *versus* h-MSCs). (H-J) Relative RQ-PCR expression of osteoprogenitor-associated genes *TNAP* (Tissue-nonspecific alkaline phosphatase, H, n=7 h-MSCs and n=3 AML-MSCs) and *OPN* (osteopontin, I, n=6 h-MSCs and n=3 AML-MSCs), and pro-inflammatory gene *PTGS2* (prostaglandin-endoperoxide synthase 2, J, n=7 h-MSCs and n=5 AML-MSCs) in h-MSCs pre-treated or not for 72 hours with Ouabain or K^+^ gluconate and in AML-MSCs. T-test was used to compare treated or AML-MSCs groups *versus* h-MSCs. All histograms show mean ± SEM; \**p*-value <0.05, \*\**p*-value <0.01, \*\*\**p*-value <0.001.

### AML blasts depolarizes MSCs V_mem_ by cell-cell contact

Considering that AML-MSCs are depolarized compared to h-MSCs, it is reasonable to speculate that AML blasts play a role in modifying V_mem_ of MSCs residing in the BM niche. We examined the V_mem_ of h-MSCs after co-culturing with AML blasts (namely induced AML-MSCs, iAML-MSCs) at different timepoints, observing that iAML-MSCs quickly increased their V_mem_, which remained constant over time reaching the depolarized V_mem_ levels of AML-MSCs (−11.8 mV, *p*<0.001, Fig. 3A-D, Supplementary Fig. 3A). To confirm that the contact between blasts and MSCs is responsible for V_mem_ changes, we demonstrated that the addition of a transwell insert in the co-culture induced a lower V_mem_ change compared to direct cell-cell contact (Supplementary Fig. 3B). We then focused on Connexin(Cx)-43-based gap junctions, fundamental in excitable tissues by facilitating transmission of electric signals between adjacent cells^28^. We previously demonstrated that Cx-43 was responsible for AML-MSCs transcriptional reprogramming^26^; here we blocked Cx-43-mediated cell-cell contact between h-MSCs and blasts by treating with a gap junction blocker, Carbenoxolone (CBX), or silencing both cell types with Cx-43 small interfering (si)RNA, and monitored V_mem_. Both strategies inhibiting cell-cell contact blocked MSCs-bioelectric shift toward a depolarized status (Fig. 3E-F). To check if AML blasts exposure was responsible for MSCs V_mem_ changes, excluding effects derived from physiological communication by heterotypic cell-cell contact, we co-cultured h-MSCs with either blasts or normal mononuclear cells (PBMCs) for 3 days, indicating that cell-cell contact alters the V_mem_ regardless of cell type, while the addition of CBX prevented this V_mem_ shift (Fig. 3G). However, only AML blasts were able to depolarize h-MSCs up to 6 days of co-culture and after blasts removal (Fig. 3H), providing the evidence that AML blasts directly induced the MSCs bioelectric shift toward depolarization.

**Fig. 3.**
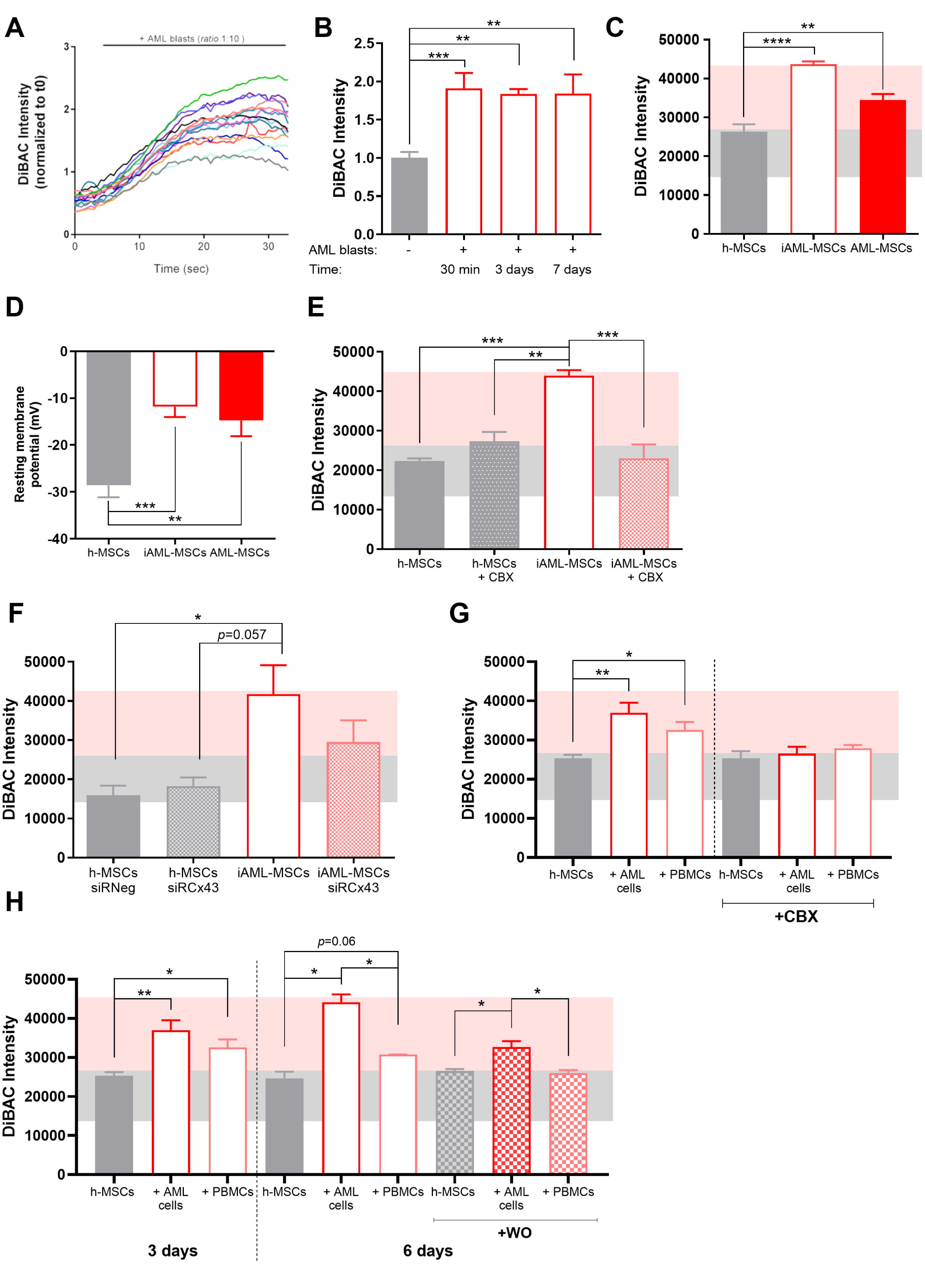
AML blasts depolarize MSCs V_mem_ by cell-cell contact. (A) Time course of DiBAC fluorescence intensity in h-MSCs during co-culture with AML blasts (ratio 1:10) calculated for the first 30 minutes of co-culture and normalized with respect to t0 fluorescence intensity value (15 cells, n=1). (B) V_mem_ variations measured by DiBAC fluorescence intensity in h-MSCs co-cultured with AML blasts for 30 min, 3 days or 7 days (n=3, t-test comparing all groups *versus* h-MSCs). DiBAC fluorescence intensity of h-MSCs cultured alone was used as control. (C) DiBAC fluorescence intensity in h-MSCs (n=9), h-MSCs cultured with AML blasts for 4 days (iAML-MSCs, n=15), and AML-MSCs (n=11). Pink and grey headbands represent the range value of AML- and h-MSCs DiBAC intensity respectively, after 4 days of culture. T-test was used to compare i-AML-MSCs or AML-MSCs *versus* h-MSCs. (D) Histogram of average resting membrane potentials (mV) measured by patch clamp in h-MSCs (n=4, 10 cells), iAML-MSCs (n=4, 10 cells) and AML-MSCs (n=5, 11 cells). T-test was used to compare i-AML-MSCs or AML-MSCs *versus* h-MSCs. (E-F) DiBAC fluorescence intensity in iAML-MSCs treated with CBX (100 µM, n=3) or silenced for Cx-43 (n=3) during co-culture with AML blasts. h-MSCs siRNeg and siRCx-43 DiBAC fluorescence intensity were used as controls. T-test was performed to compare h-MSCs or i-AML-MSCs treated with CBX/SirCx43 *versus* i-AML-MSCs. (G) Fluorescence intensity measured by DiBAC staining after 3 days of h-MSCs co-culture with AML primary cells or normal PBMCs (n=7), with or without the addition of CBX (n=3). T-test was performed to compare all groups with their respective h-MSCs group. (H) DiBAC intensity of h-MSCs co-culture with AML primary cells or normal PBMCs for 3 days (n=7), 6 days (n=3), or 3 days of co-culture followed by 3 days of washout (WO, n=3). T-test used to compare all groups with their respective h-MSCs group. All histograms show mean ± SEM; \**p*-value <0.05, \*\**p*-value <0.01, \*\*\**p*-value <0.001, \*\*\*\**p*-value <0.0001.

### MSCs V_mem_ pharmacological hyperpolarization restores healthy features

Considering that h-MSCs change their bioelectricity when leukemia occurs, we investigated whether depolarized AML-MSCs could recover healthy hyperpolarized V_mem_ and healthy features. We hyperpolarized AML-MSCs V_mem_ by using Lubi and IVM and, after reducing V_mem_ (Fig. 1J), we observed a lowered proliferation rate similar to h-MSCs (IVM: −1.9-fold, *p*<0.05, Fig. 4A). Moreover, hyperpolarization reduced 32D cells support when cultured without IL-3 (*p*<0.01, Fig. 4B-C), and recovered the anti-inflammatory potential, confirmed by a greater anti-angiogenic activity and higher potential to suppress T-cell activation (*p*<0.05, Fig. 4D-E) at the same strength of h-MSCs. Moreover, we observed a significantly reduced expression of osteo-related genes *TNAP* (*p*<0.0001, Fig.4F) and *OPN* (*p*<0.05, Fig. 4G), recovering h-MSCs level.

**Fig. 4.**
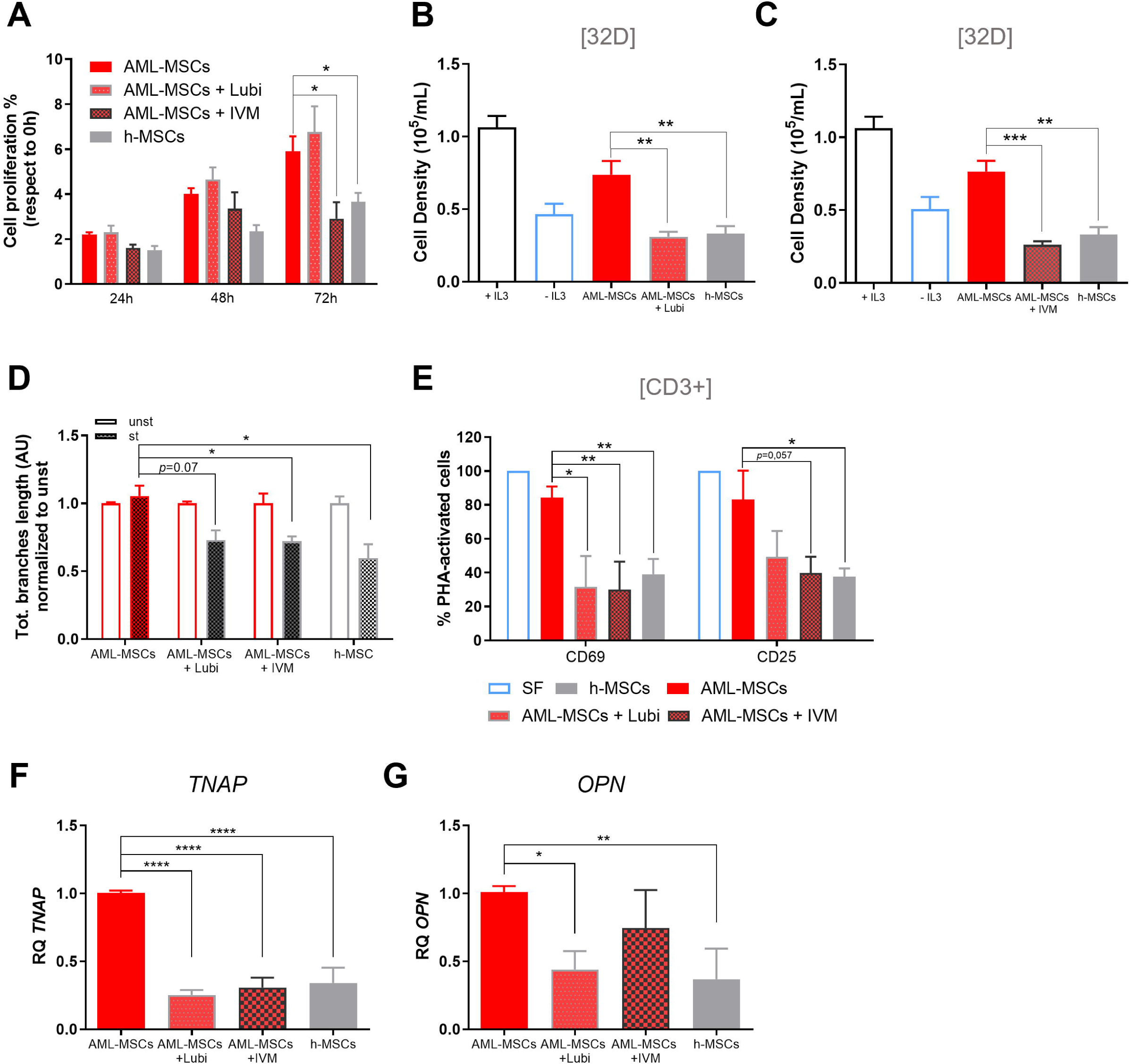
AML-MSCs V_mem_ pharmacological hyperpolarization restore a healthy phenotype. (A) Cell proliferation measured by Presto Blue assay in h-MSCs and AML-MSCs treated or not with 10 µM Lubi or 1 µM IVM for 72 hours (n=4, t-test comparing all groups *versus* AML-MSCs). (B-C) Cell density of murine IL-3–dependent 32D cell line cultured in the presence or absence of IL-3, with h-MSCs, and compared with 32D cell line co-cultured on AML-MSCs layer treated with Lubi (B, n=5) or IVM (C, n=5). T-test was performed to compare treated groups or h-MSCs *versus* AML-MSCs, with ±IL3 used as experimental control. (D) Total branches length of HUVEC tubes by using conditioned medium derived from h-MSCs (n=4) and AML-MSCs untreated (n=7) or pre-treated with Lubi (n=8) or IVM (n=5) for 72 hours and then stimulated (st) or not (unst) with a pro-inflammatory cytokine cocktail for 24 hours; tube formation was evaluated after 4 hours and normalized to unst condition (AU: arbitrary unit, t-test comparing treated or h-MSCs stimulated groups *versus* stimulated AML-MSCs). (E) Percentage of PHA-stimulated CD3^+^ T cells expressing CD69 and CD25 after 72 hours of co-culture on a layer of h-MSCs or AML-MSCs treated or not with Lubi (10 µM) or IVM (1 µM), relative to SF condition (n=4, t-test comparing treated or h-MSCs groups *versus* AML-MSCs). (F-G) Relative expression of osteoprogenitor-associated genes *TNAP* (F, n=7 AML-MSCs, n=4 h-MSCs) and *OPN* (G, n=6 AML-MSCs, n=4 h-MSCs) in h-MSCs and AML-MSCs treated or not for 72 hours with Lubi (10 µM) or IVM (1 µM) and measured by RQ-PCR (t-test comparing treated or h-MSCs groups *versus* AML-MSCs). All histograms show mean ± SEM; \**p*-value <0.05, \*\**p*-value <0.01, \*\*\**p*-value <0.001, \*\*\*\**p*-value <0.0001.

To definitively exclude a role of the drugs *per se*, we treated h-MSCs with hyperpolarizing drugs (IVM, Lubi) and AML-MSCs with depolarizing agents (K^+^ gluconate, Ouabain) showing that neither cell proliferation nor immunomodulatory functions were affected (Supplementary Fig. 4A-B), thus excluding direct effects of the drugs on these functions.

### CaV1.2 expression is linked to V_mem_

Since we established a link among blasts, MSCs and depolarization, we investigated whether Ca^2+^ and CaV1.2 channel were involved. We previously documented a different expression of CaV1.2 between h- and AML-MSCs^26^. Here, we showed that h-MSCs reduced CaV1.2 protein expression when co-cultured with AML blasts (<44%, *p*<0.05, Fig. 5A, Supplementary Fig. 5). We observed a similar down-regulation of CaV1.2 expression in h-MSCs depolarized for 72 hours with Ouabain or K^+^ gluconate (<35%, *p*<0.05, Fig. 5A). In line with this finding, a 72-hour treatment of AML-MSCs with hyperpolarizing agents showed an up-regulation of CaV1.2 protein expression (1.8-fold, *p*<0.01, Fig. 5B, Supplementary Fig. 5). To corroborate these findings, we blocked CaV1.2 with Lercanidipine, a selective CaV1.2 channel blocker previously shown to induce cell death in AML-MSCs (expressing low CaV1.2 levels), whereas not altering Ca^2+^ influx nor viability of h-MSCs (expressing higher levels of CaV1.2)^26^. In detail, we pharmacologically depolarized h-MSCs or hyperpolarized AML-MSCs, and after 72 hours we treated cells with Lercanidipine, showing that h-MSCs, when depolarized, became sensitive to Lercanidipine reducing their viability similarly to AML-MSCs (*p*<0.05, Fig. 5C); on the contrary, by hyperpolarizing AML-MSCs, cells increased CaV1.2 expression and did not respond to lercanidipine similarly to h-MSCs (Fig. 5D).

**Fig. 5.**
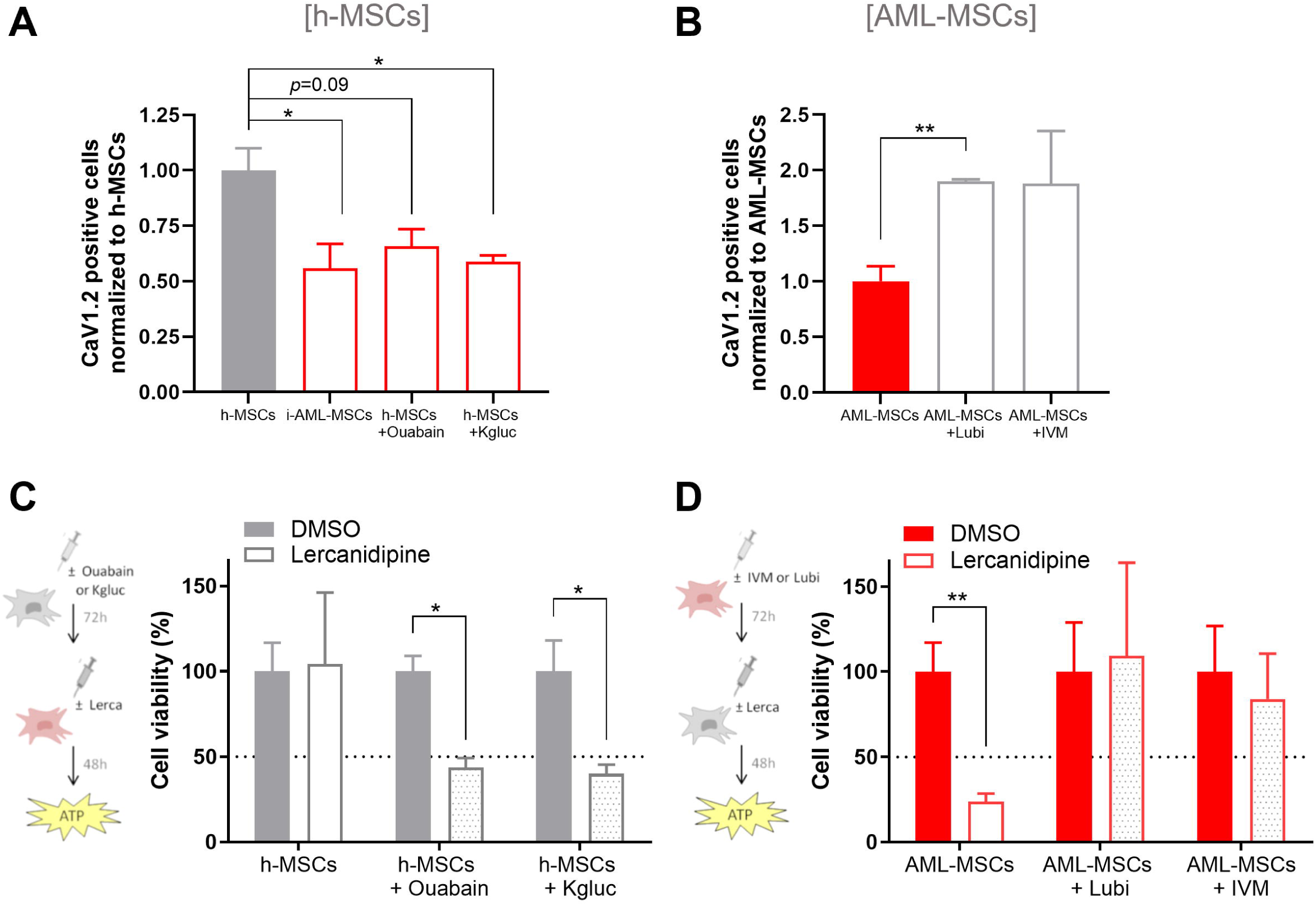
V_mem_ of MSCs controls CaV1.2 expression. (A) CaV1.2 expression in h-MSCs when co-cultured with AML cells (red bar) for 4 days or treated with Ouabain or K^+^ gluconate (red bar) for 72 hours, relative to h-MSCs (grey bar), analyzed by flow cytometry (n=3, t-test comparing all groups *versus* h-MSCs). (B) CaV1.2 expression in AML-MSCs when treated with Lubi or IVM (grey bar, n=4) for 72 hours, normalized to AML-MSCs (red bar), analyzed by flow cytometry (t-test comparing all groups *versus* AML-MSCs). (C) Cell viability analyzed by ATP of h-MSCs pre-treated or not for 72 hours with Ouabain or K^+^ gluconate and then incubated for 48 hours with lercanidipine. ATP values were normalized to their respective untreated control (DMSO, n=6, t-test comparing all groups *versus* untreated h-MSCs). (D) Cell viability, measured by ATP, of AML-MSCs pre-treated or not for 72 hours with Lubi or IVM and then for 48 hours with lercanidipine, relative to DMSO value (n=8, t-test comparing all groups *versus* untreated AML-MSCs). Dotted line represents 50% cell viability. All histograms show mean ± SEM; \**p*-value <0.05, \*\**p*-value <0.01.

### CaV1.2 over-expression reverts AML-MSCs phenocopying h-MSCs properties, including hematopoietic stem cells support

Considering the lower CaV1.2 expression in AML-MSCs with respect to h-MSCs, and the correlation of its levels with V_mem_, we hypothesized that recovering CaV1.2 expression in AML-MSCs could correct their function. We first generated a plasmid containing CaV1.2 and tested its functionality on HEK293T, observing that CaV1.2 over-expression (Supplementary Fig. 6A) led to an increased intracellular Ca^2+^ uptake in response to KCl depolarization (Supplementary Fig. 6B-C). Next, we produced a CaV1.2 third-generation lentiviral vector (LV) for a gene therapy approach in AML-MSCs. Fourteen days post-infection, we assessed CaV1.2 mRNA levels by RQ-PCR confirming its over-expression in transduced AML-MSCs, namely reverted AML-MSCs (re-AML-MSCs) (*p*<0.05, Fig. 6A), further confirmed by immunofluorescence staining of CaV1.2 (Fig.6B). Therefore, we measured re-AML-MSCs V_mem_ observing a more hyperpolarized status, reaching the DiBAC intensity of h-MSCs (grey headband, *p*<0.001, Fig. 6C). However, despite the transduction efficiency (mCherry^+^) ranging from 10% to 45%, we measured a more hyperpolarized V_mem_ also in mCherry-negative cells with respect to un-transduced (UT) MSCs, this latter finding arguing that the recovered hyperpolarized V_mem_ propagated to neighboring AML-MSCs. To further elucidate this finding, we mixed AML-MSCs UT with re-AML-MSCs (ratio 1:1) and demonstrated that V_mem_ was hyperpolarized in all cells and resembled that of h-MSCs, supporting the hypothesis that re-AML-MSCs influenced the surrounding AML-MSCs V_mem_ (Fig. 6C). We performed gene expression profile analysis, and PCA showed that h- and re-AML-MSCs clustered together but separately from AML-MSCs (Fig. 6D), suggesting that CaV1.2 over-expression triggered AML-MSCs to hyperpolarization and to a healthy gene expression reprogramming. We also investigated how re-AML-MSCs were able to propagate the depolarization wave to neighbor cells. We previously demonstrated in a 3D niche that MSCs contact local and distant cells by gap junctions and tunneling nanotubes (TNTs), thin F-actin-based membranous protrusions of MSCs, enabling the communication with the cytoplasm of neighboring cells^26^. We found that re-AML-MSCs formed TNTs with surrounding cells and that CaV1.2 localized and transited along these structures (Fig.6E). Moreover, treatment with CBX impaired the formation of TNTs between re-AML-MSCs and AML-MSCs, significantly preventing the achievement of the bioelectric state of re-AML-MSCs in AML-MSCs (Fig. 6F).

**Fig. 6.**
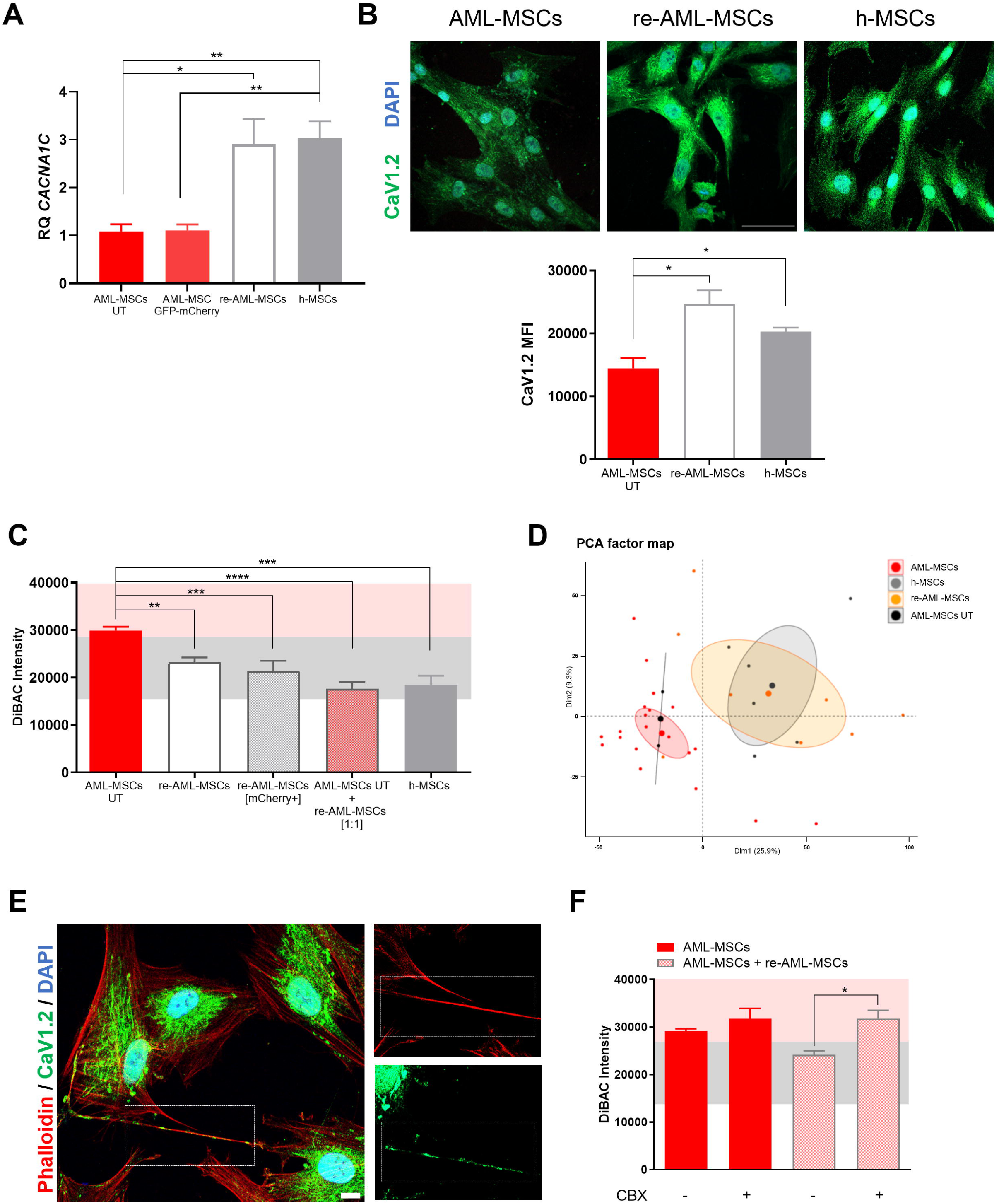
CaV1.2 expression controls MSCs V_mem_ depolarization. (A) Relative *CACNA1C* (CaV1.2) mRNA expression measured by RQ-PCR in AML-MSCs UT, transduced with control GFP-mCherry or CaV-mCherry LVs (re-AML-MSCs, n=11), or h-MSCs (n=3, t-test comparing all groups *versus* AML-MSCs). (B) Representative images of immunofluorescence staining of CaV1.2 (green) and DAPI nuclear counterstain (blue) and relative quantification of CaV1.2 mean fluorescence intensity (MFI) (20X, 20 cells/image, 5 images, scale bar=50 µm, n=3 AML-MSCs and re-AML-MSCs, n=2 h-MSCs, t-test comparing all groups). (C) DiBAC intensity of AML-MSCs (n=9), re-AML-MSCs total (n=9) or gated on mCherry positive cells (n=6), in a mixed culture of re-AML-MSCs and AML-MSCs UT (ratio 1:1, n=4), and h-MSCs as control (n=3). T-test was adopted to compare all groups *versus* AML-MSCs). (D) Principal Component Analysis (PCA) performed on gene expression data of h-MSCs (grey, n=6), AML-MSCs (red, n=21), AML-MSCs UT (black, n=2), and re-AML-MSCs (orange, n=7). Big dots represent the cluster centroid and dots represent samples. (E) Representative images showing the presence of CaV1.2 channel inside the TNTs. Green indicates CaV1.2, blue indicates nuclei staining with DAPI, and red indicates F-actin staining with Alexa647-Phalloidin (100X, scale bar=10 μm). (F) DiBAC intensity in AML-MSCs and in a mixed culture of re-AML-MSCs and AML-MSCs UT (ratio 1:1), treated or not with 100 µM Carbenoxolone (CBX) for 72 hours (n=6). T-test was applied to compare treated MSCs *versus* untreated. All histograms show mean ± SEM; \**p*-value <0.05, \*\**p*-value <0.01, \*\*\**p*-value <0.001, \*\*\*\**p*-value <0.0001.

Considering that re-AML-MSCs are reprogrammed to h-MSCs, we explored re-AML-MSCs functions. We documented that re-AML-MSCs reduced the growth of 32D cell line in the absence of IL-3 with respect to AML-MSCs (*p*<0.05, Supplementary Fig. 7A) and rescued immunosuppressive properties (Supplementary Fig. 7B); however, proliferation and anti-inflammatory potential of re-AML-MSCs were not significantly affected with respect to AML-MSCs after 72 hours (Supplementary Fig. 7C-D). To further corroborate re-AML-MSCs reprogramming, we used our previously established 3D BM niche model^26^ and measured MSCs V_mem_ in a more complex *in vitro* system, confirming that AML-MSCs were depolarized with respect to h-MSCs and re-AML-MSCs, that were hyperpolarized (Fig. 7A). Therefore, considering the impact of AML-MSCs in sustaining AML blasts (Supplementary Fig. 8A), we evaluated blasts proliferation in 3D when co-cultured with different MSCs. Results showed a reduction of AML cells proliferation when co-cultured with h-MSCs or re-AML-MSCs with respect to AML-MSCs or h-MSCs depolarized with K^+^ gluconate (Fig. 7B), this supporting the direct role of MSCs in sustaining TME through V_mem_. Since h-MSCs play a key role in controlling HSCs homeostasis and functions, we investigated re-AML-MSCs support to normal HSCs (CD34^+^). *CXCL12* and *ANGPT1* expression, critical for HSCs engraftment, homing and proliferation^29,30^, was found remarkably up-regulated in re-AML-MSCs compared with AML-MSCs. *VEGFA*, which regulates HSCs quiescence^31^, was significantly down-regulated in re-AML-MSCs (*p*<0.001, Supplementary Fig. 9A). Hence, we asked whether the recovery of healthy features in re-AML-MSCs could restore their hematopoietic supportive capacity *in vitro* and *in vivo*. We compared the expansion of CD34^+^ cells cultured alone (SF) or on a layer of h-, AML, or re-AML-MSCs, confirming that AML-MSCs decreased HSCs proliferation (−2.2-fold, *p*<0.001, Fig. 7C), whereas re-AML-MSCs and h-MSCs supported CD34^+^ proliferation (*p*<0.001, Fig. 7C), also validated by Ki-67 positivity (Fig. 7D). *In vivo*, CD34^+^ HSCs were co-infused by tail vein injection into sub-lethally irradiated NOD/SCID/IL-2Rγ null (NSG) mice together with h-, AML-, or re-AML-MSCs. For 10 weeks after transplantation, peripheral blood (PB) samples were analyzed for the presence of human CD45^+^ (hCD45^+^) cells. In line with *in vitro* data, we detected a significantly lower percentage of hCD45^+^ in the PB of NSG mice receiving CD34^+^ cells in co-transplantation with AML-MSCs as compared to re-AML-MSCs (n=6 mice per group, *p*<0.05, Fig. 7E), as well as h-MSCs. Since there are debated results on the role of MSCs in improving hematopoietic reconstitution^32,33^, we strengthened re-AML-MSCs capability of sustaining engraftment of CD34^+^ cells by using a second mice strain (humanized NOG-EXL, Supplementary Fig. 9B), confirming these findings. Thus, to link the increased CD34^+^ engraftment to the presence of MSCs, we monitored MSCs *in vivo*. In detail, we injected mCherry-transduced MSCs *in vivo* and sacrificed mice at 24, 48, and 72 hours post-injection, demonstrating, in the window of CD34^+^ homing to murine BM, that MSCs were increasingly repopulating the BM, and migrated also in lung, spleen and liver, as expected^34^ (Supplementary Fig. 9C). Then, CD34^+^ cells were pre-stained with CFSE to test their engraftment and proliferation *in vivo* when co-injected with h-, AML- or re-AML-MSCs. Results showed that re-AML-MSCs improved CD34^+^ cell proliferation with respect to AML-MSCs (reduction of %CFSE^+^ cells in BM, Supplementary Fig. 10A-B). These results documented that, for three days after injection, MSCs reside in the BM playing a direct role in contributing to HSCs engraftment and proliferation, thus sustaining hematopoiesis as demonstrated by hCD45^+^ cells in mice PB from 5 to 10 weeks post CD34^+^ and h- or re-AML-MSCs co-injection (Fig. 7E, Supplementary Fig.9B). These data highlight that MSCs co-transplantation with donor’s HSCs have the potential to increase an early graft, as previously reported^34–36^. Altogether, autologous re-AML-MSCs promote a successful early HSCs anchorage in the BM and contribute to their proliferation and to hematopoiesis recovery *in vivo*.

**Fig. 7.**
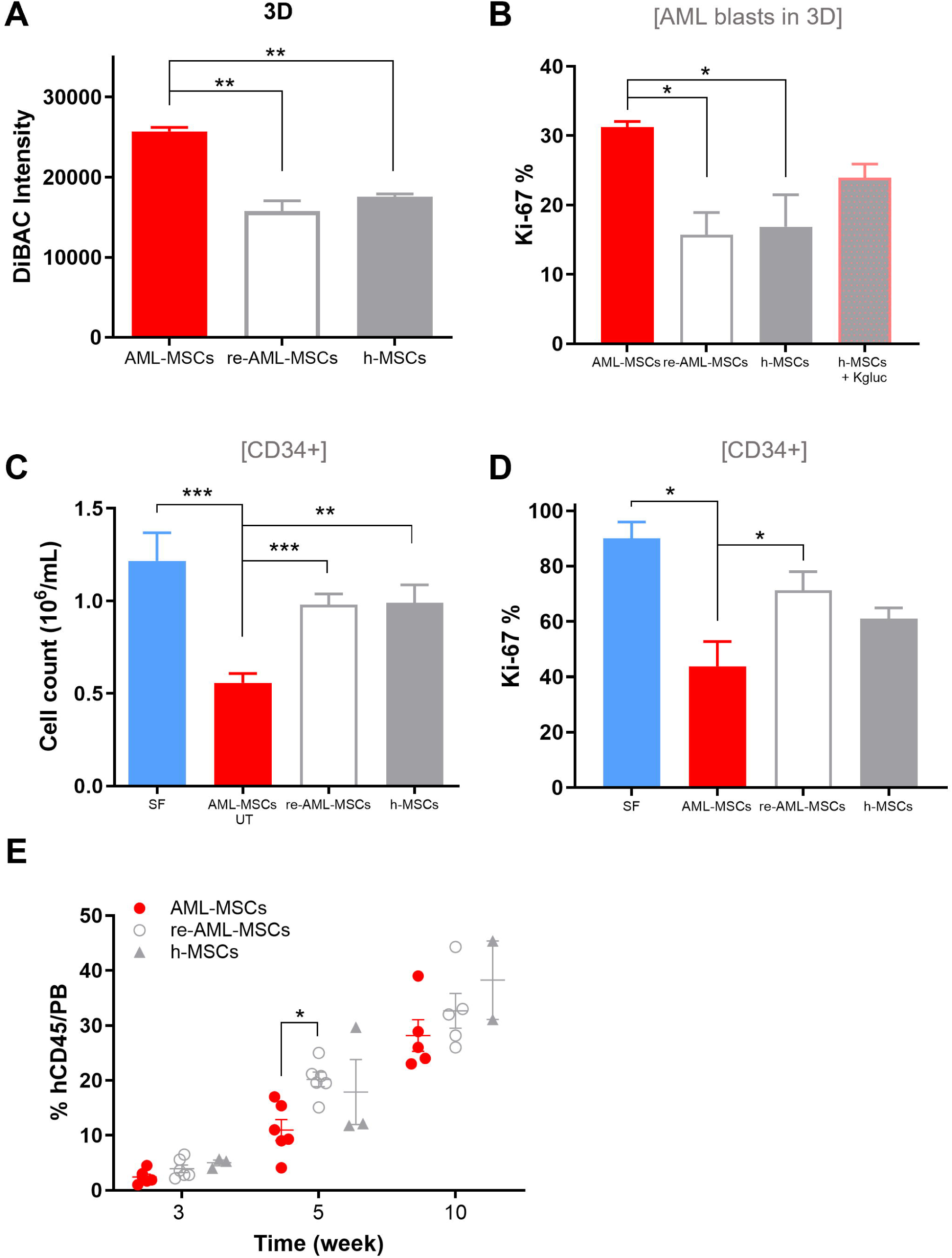
V_mem_ of MSCs influences leukemia proliferation and normal hematopoiesis. (A) DiBAC fluorescence intensity of h-MSCs (n=2), AML-MSCs (n=3) and re-AML-MSCs (n=2) in 3D. (B) Percentage of Ki-67 positive AML blasts after a 7-day co-culture in 3D with AML-MSCs (n=3), re-AML-MSCs (n=2), h-MSCs treated or not with K^+^ gluconate (n=2, t-test comparing all groups). (C) Cell density of CD34^+^ cells after a 5-day co-culture with AML-MSCs UT, re-AML-MSCs (n=9) and h-MSCs (n=3). CD34^+^ cells cultured in SF condition were used as control. (D) Percentage of Ki-67 positive CD34^+^ cells after 5 days of co-culture with AML-MSCs UT, re-AML-MSCs and h-MSCs (n=3). (E) Percentage of human CD45 cells in NSG mice PB at 3, 5, and 10 weeks after tail vein co-infusion of 1.0×10^5^ human CB CD34^+^ cells with 1.0×10^6^ h-MSCs (n=3 mice), AML-MSCs (n=6 mice), or re-AML-MSCs (n=6 mice, t-test comparing all groups). All histograms show mean ± SEM; \**p*-value <0.05, \*\**p*-value <0.01, \*\*\**p*-value <0.001.

## CONCLUSIONS

Bioelectric signaling emerged in the last decades as an important controller of cell growth, cell migration, tumor progression, metastasis, and TME^11^. It is nowadays accepted that aberrant bioelectrical interactions, mediated by altered ion channels expression and activity, can promote abnormal depolarization of resting potential and cell signaling also in *non*-excitable cells^14,20^. MSCs are a biologically relevant component of the BM niche, being able to support hematopoiesis but also to differentiate along mesenchymal and non-mesenchymal *lineages* to acquire immunomodulatory, reparative, and anti-inflammatory properties^37–39^. Recently, the therapeutic potential of MSCs has expanded beyond conventional applications, incorporating innovative cell-based strategies such as engineered MSCs and CAR-MSCs. These approaches aim to enhance the efficacy of cancer therapy by leveraging MSCs tumor-homing properties and modulating the tumor microenvironment, broadening their role in targeted cellular therapies^40–43^. We previously characterized MSCs derived from AML patients, demonstrating that lercanidipine, a CaV1.2 channel blocker selective for AML-MSCs, impairs leukemia progression^26^. CaV1.2 is highly expressed in the brain and peripheral nervous system regulating neuropathic pain^44^, but its role in MSCs is largely unexplored. Here we found that Ca^2+^ dynamics was different in h- and AML-MSCs, and questioned if tumor-specific variations in ion balance contributes to confer leukemia aggressiveness, and induce the TME by affecting MSCs functions. We found that decreased density of CaV1.2 in AML-MSCs mediated an aberrant Ca^2+^ uptake and membrane depolarization with respect to h-MSCs which are hyperpolarized, also confirmed in a 3D model of the BM niche. These findings indicate that AML-MSCs help shape the leukemic niche through their electrical properties^8,21^. In line with this hypothesis, co-culture experiments of h-MSCs and AML blasts revealed that a stable MSCs depolarization occurred upon blasts exposure, reaching a depolarized V_mem_ comparable to AML-MSCs. Particularly, MSCs retained a depolarized V_mem_ only after contact with blasts, whereas their V_mem_ returned to basal levels following PBMCs removal, this providing evidence that AML blasts directly induce the irreversible MSCs bioelectric shift toward depolarization. So far, the underlying mechanism responsible for the propagation of the bioelectric state from tumor cells to neighbor stromal cells is largely underexplored. We previously observed that AML cells and AML-MSCs formed networks through gap junctions in 3D^26^. Here, we showed that the interruption of blast-MSC communication, inhibiting or silencing Cx-43, prevented the achievement of the depolarization status characterizing AML-MSCs, demonstrating that AML blasts drive MSCs bioelectric and transcriptomic transition by cell-cell contact. Interestingly, in the leukemia context, recent reports showed that disruption of Cx-43 on MSCs influences HSCs homeostasis and leukemia proliferation, as well as chemoresistance^45,46^.

Since ion channels transmit extensive information patterns between different cell types and the environment, we hypothesized that tumor cells may not depend exclusively on altered genes for progression, but also on non-autonomous cell interactions and ion balance. By pharmacological modulation of ion channels activity, we revealed that V_mem_ depolarization induced a pro-leukemia-like phenotype on h-MSCs, supporting AML proliferation in a 3D model. Conversely, hyperpolarizing AML-MSCs suppressed their pro-leukemic phenotype, suggesting that V_mem_ hyperpolarization alone, independently of ion-specific effects or the scaffolding roles of ion channel proteins, can modulate cell functions in the absence of other molecular interventions (such as gene knockout or RNA interference). Likewise, even if ion channel dysregulation in spatial or temporal tumor heterogeneity remains to be further explored, we lay the basis for considering bioelectricity as a potential feature influencing treatment efficacy, clonal evolution and disease progression^47^. In line with these findings, it was crucial to confirm the role of electrical signals in MSCs. We demonstrated that CaV1.2 over-expression by lentiviral vectors in reverted AML-MSCs (re-AML-MSCs) could, itself, repolarize AML-MSCs toward a healthy hyperpolarized state, that propagated in a wave typical of excitable cells, and restored the h-MSCs gene expression profile. We rationalized how electrical signals propagate and showed a nanotubes-mediated CaV1.2 trafficking that caused V_mem_ hyperpolarization in neighbor cells. This latter finding of CaV1.2 hijacking from re-AML-MSCs to AML-MSCs emerged as a novel mechanism to restore healthy MSCs. In addition, our gene therapy approach significantly supported hematopoiesis by improving the survival and expansion of HSCs (CD34^+^) *in vitro* and *in vivo*. HSCs homing and engraftment still represent an important issue leading to graft failure. Several works revealed that the presence of a functional BM microenvironment capable of sustaining HSCs engraftment, expansion and differentiation, and reducing graft-versus-host disease, is a fundamental requisite for a successful transplantation outcome^48–50^. Here, to face an important therapeutic issue related to MSCs limited homing and long-term persistence to target organs following administration, we tracked the presence of MSCs with HSCs *in vivo* three days after injection, observing a remarkable MSCs capacity to sustain HSCs homing and proliferation, confirming a direct role of MSCs in HSCs engraftment. Moreover, CaV1.2 restoration to autologous AML-MSCs repolarized and restored Ca^2+^ dynamics and a healthy MSCs program. Despite the many questions that still need to be answered, these results of a gene therapy approach on MSCs in the context of HSCT has an enormous potential^32,34–36^, as supported by several different clinical trials recently emerged (NCT01045382, NCT06171906, NCT02291770, NCT02379442), including CAR-MSCs which represent a widely applicable therapeutic technology in the field of immune-therapy^40^.

In conclusion, our findings reveal a groundbreaking mechanism by which CaV1.2 expression modulates Ca²[ levels, influencing MSCs V_mem_ through a dynamic feedback loop dependent on cell-cell interactions. These discoveries unveil an unprecedently recognized bioelectric code governing TME in leukemia, with MSCs V_mem_ as a key regulatory feature in BM niche remodeling and in hematopoietic support, thus marking a major step forward in understanding leukemogenesis. Overall, harnessing bioelectricity to restore homeostasis in leukemia could redefine future approaches in precision medicine, offering new hope for novel incoming cellular interventions.

## ACKNOWLEDGEMENTS

We thank the staff of the laboratory of the Division of Pediatric Hematology, Oncology and Stem Cell Transplant, Women’s and Children’s Health Department for diagnostic activity and biobanking (Biobanca Oncologia Pediatrica-BBOP). The authors are grateful to all AIEOP centers. We thank Matteo Barioni and Barbara Sartini of the Cord Blood Bank of the Division of Pediatric Hematology, Oncology and Stem Cell Transplant, at Women’s and Children’s Health Department of the University-Hospital of Padova. Giulia Borella and Maddalena Benetton were supported by an AIRC fellowship for Italy.

## RESOURCE AVAILABILITY

All original data produced in this study are available upon request to the lead contact at.

The datasets generated during the current study are available in the GEO repository, with the access code GSE248681. To perform the analyses reported in this study, data were analyzed together with those previously deposited at GSE169428 (Borella G. *et al.*, Blood 2021 Aug 19;138(7):557-570).

## AUTHOR CONTRIBUTIONS

G.Bore., M.Be, A.D.R. and M.P. contributed to conceptualization. G.Bore., M.Be., A.D.R., G.L., G.Bori., A.C., D.L.P., C.T., M.Bo., S.B. and M.P. contributed to methodology. G.Bore., M.Be., A.D.R., D.L.P. and S.B. contributed to validation. G.Bore., M.Be., A.D.R., G.Bori., D.L.P. and S.B. contributed to formal analysis. G.Bore., M.Be., A.D.R., G.L., G.Bori., A.C., D.L.P. and C.T. contributed to investigation. G.Bore., M.Be., A.D.R., G.Bori., D.L.P., M.Bo., S.B. and M.P. were involved in data curation, and S.B. performed software analyses. G.Bore., M.Be., A.D.R. contributed to visualization. M.Bo., F.L. and M.P. provided study samples and materials. G.Bore., M.Be., A.D.R. and M.P. wrote the original draft, and G.Bore., M.Be., A.D.R., C.T., F.L., M.P. reviewed and edited the manuscript. C.T. and M.P. administered the project, supervised by F.L. and M.P.. M.P. acquired fundings to support this study. All authors read and approved the final manuscript.

