## Supplemental Figures for "Leukemic Cells Manipulate MSCs Bioelectrical Signals to Reshape the Bone Marrow Niche"

Borella G. *et al.*

### SUPPLEMENTARY FIGURES.

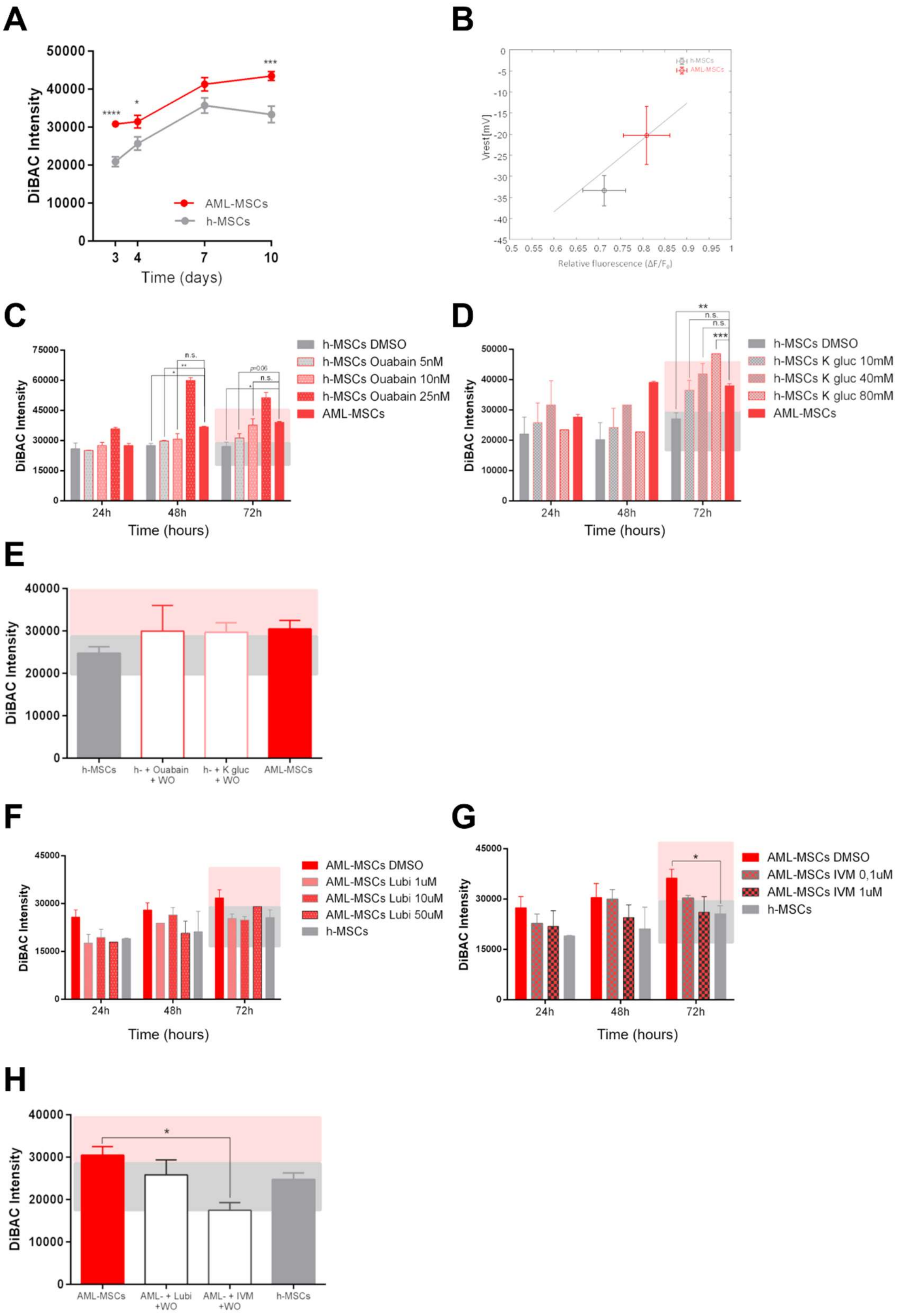

**Supplementary Fig 1. MSCs $V_{mem}$ and its modulation by drugs.** (A) Time course of fluorescence intensity calculated from DiBAC fluorescence staining in AML-MSCs (n=14) and h-MSCs (n=7) 3, 4, 7 and 10 days after seeding. Graph shows the mean ± SEM. (B) Linear regression model between mean relative fluorescence intensity ($\Delta F/F_0$) of DiBAC and Vrest (mV) values derived from patch clamp calculated for each group ($R^2$=0.69, n=5 AML-MSCs, n=4 h-MSCs). (C) Dose-time curve of DiBAC fluorescence intensity in h-MSCs treated with Ouabain at 5, 10, 25 nM for 72 hours (n=5). AML-MSCs were used as control (n=3). (D) Dose-time curve of DiBAC fluorescence intensity in h-MSCs treated with $K^+$ gluconate at 10, 40, 80 mM for 72 hours (n=4). AML-MSCs were used as control (n=3). (E) Fluorescence intensity calculated from DiBAC fluorescence staining of h-MSCs (n=3) incubated with 10 nM Ouabain or 40 mM $K^+$ gluconate for 3 days followed by incubation with control medium (Washout, WO) for additional 3 days respect to untreated-AML-MSCs (n=2). (F) Dose-time curve of DiBAC fluorescence intensity in AML-MSCs treated with Lubiprostone at 1, 10, 50 µM for 72 hours (n=3). Healthy-MSCs were used as control (n=3). (G) Dose-time curve of DiBAC fluorescence intensity in AML-MSCs treated with IVM at 0,1 and 1 µM for 72 hours (n=3). Healthy-MSCs were used as control (n=3). (H) DiBAC fluorescence intensity of AML-MSCs (n=3) incubated with 10 µM Lubiprostone or 1 µM IVM for 3 days followed by incubation with control medium (WO) for additional 3 days. Healthy-MSCs were used as control (n=3).

All histograms show mean ± SEM; **p*-value <0.05, ***p*-value <0.01, ****p*-value <0.001, ****p*-value <0.001.

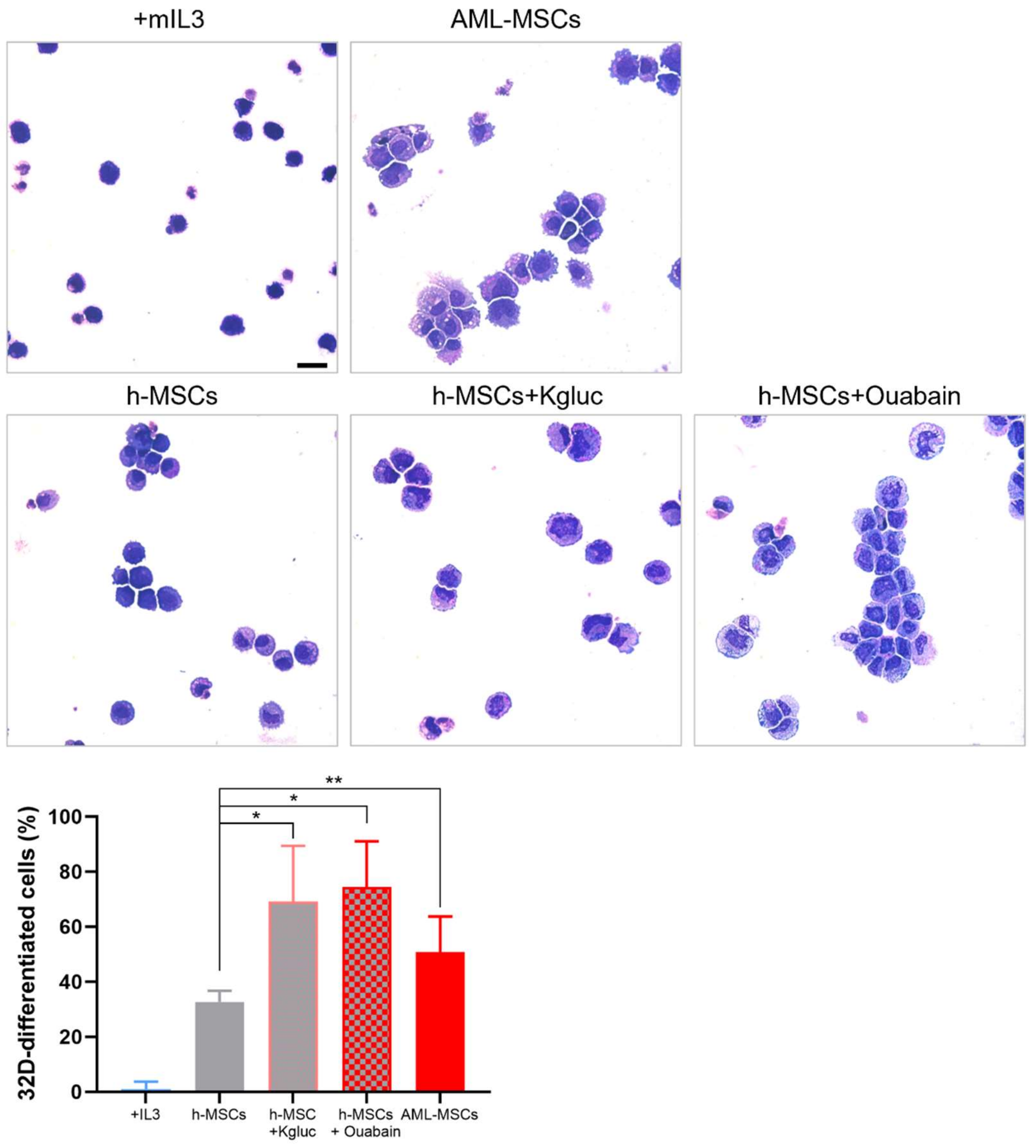


**Supplementary Fig. 2. MSCs role in 32D cell differentiation.** Upper panel: Representative May-Grünwald Giemsa staining of 32D after a 72-hour co-culture with AML-MSCs, h-MSCs treated or not with K+ gluconate or Ouabain. Scale bar=10 µm. Lower panel: histogram reporting the percentage of differentiated 32D cells (n=5 pictures per condition, minimum 100 cells analyzed). Murine IL3 was used as experimental control.

The histogram shows mean ± SEM; **p*-value <0.05, ***p*-value <0.01.

A

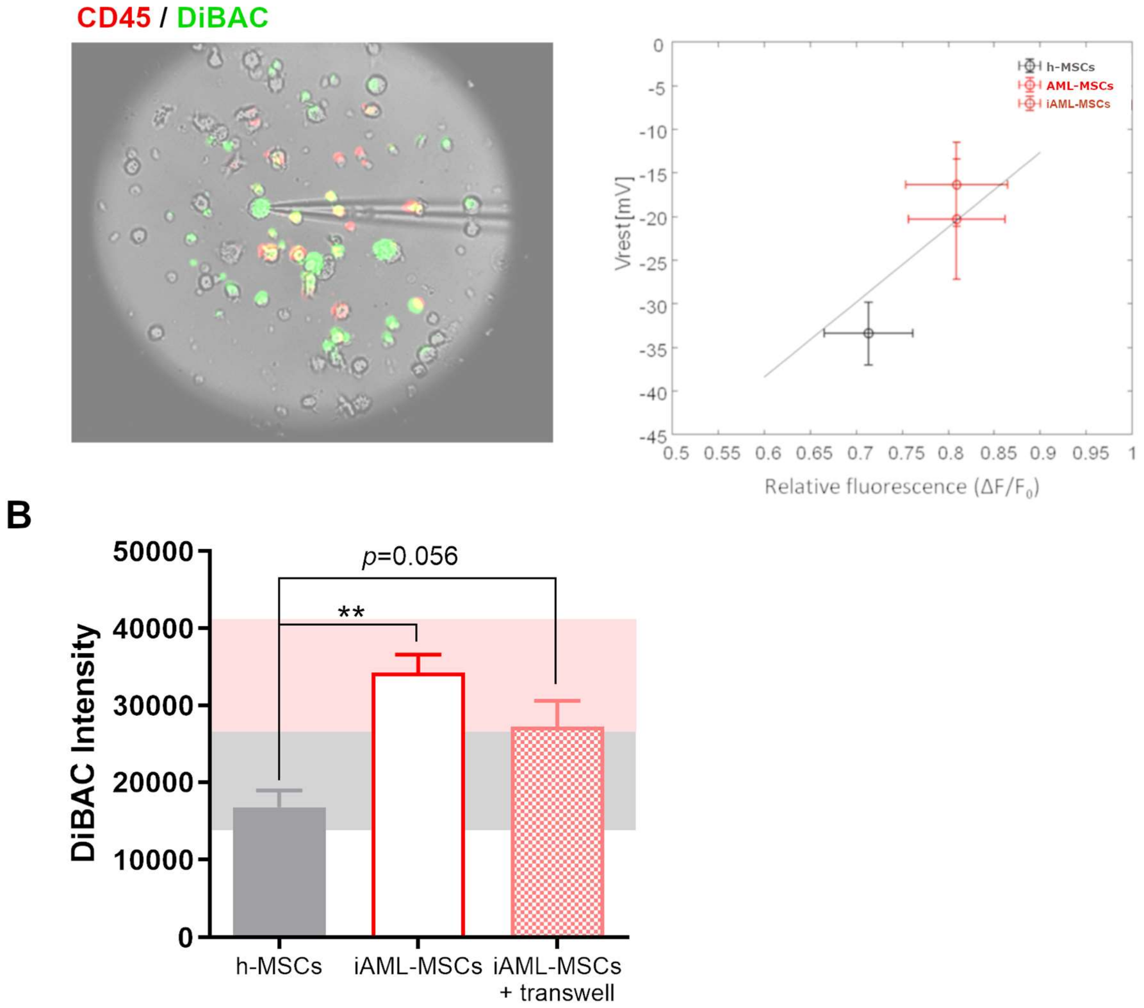


**Supplementary Fig. 3. AML blasts depolarize MSCs $V_{mem}$.** (A) Representative image of patch clamp technique on co-culture dish with AML blasts ($CD45^+$, red) and MSCs loaded with DiBAC (green) (right panel) and linear regression model of mean fluorescence intensity $\Delta F/F_0$ of DiBAC and Vrest (mV) values obtained by patch clamp calculated for each group (left panel). (B) DiBAC $V_{mem}$ measure of iAML-MSCs when a transwell insert was added during the co-culture with AML blasts (n=5, t-test comparing all groups *versus* h-MSCs). Healthy-MSCs were used as control.

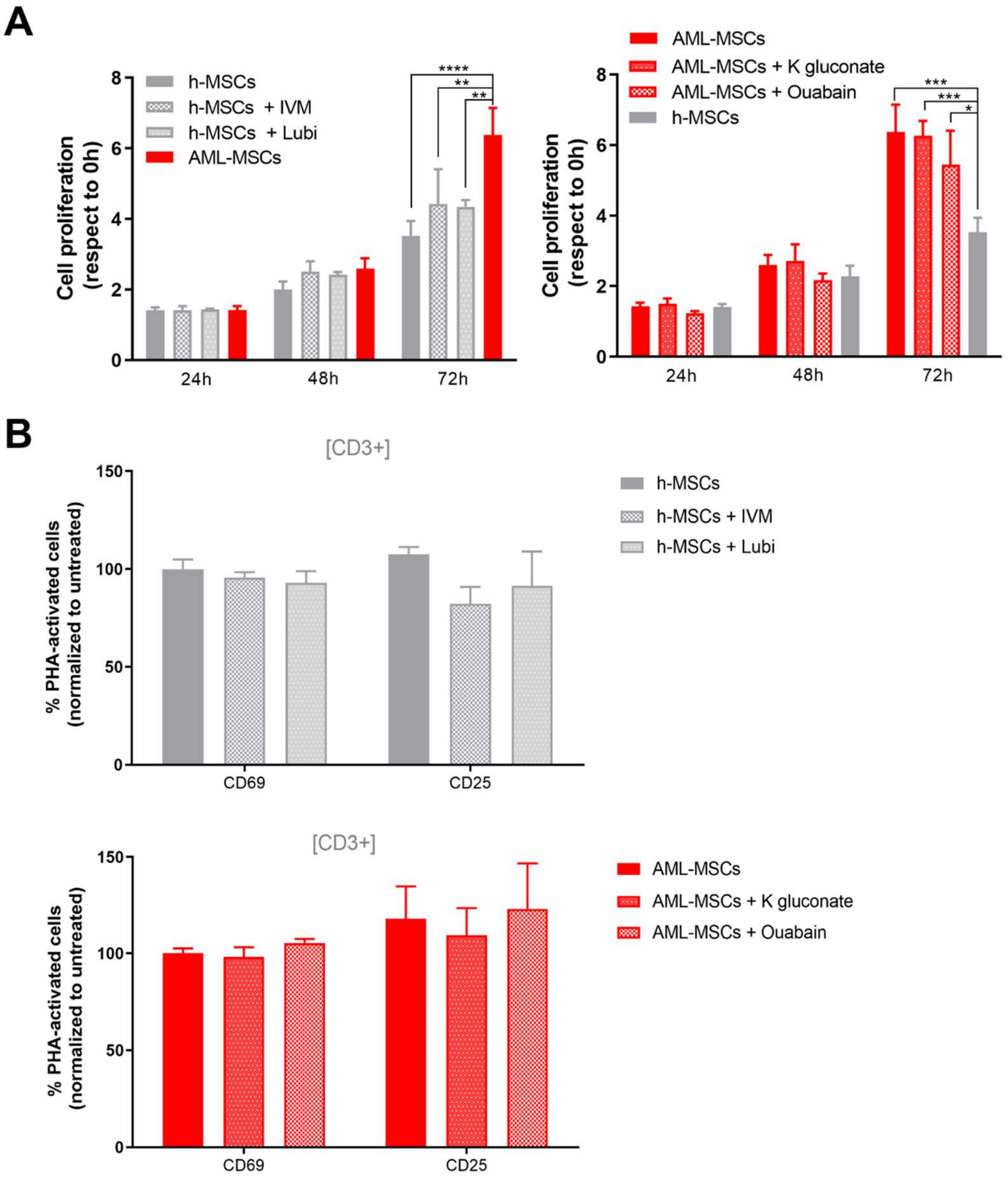


**Supplementary Fig. 4. Drug effects in MSCs.** (A) Cell proliferation of h-MSCs, AML-MSCs, h-MSCs treated with 10 μM Lubi or 1 μM IVM and AML-MSCs treated with 40 mM $K^+$ gluconate or 10 nM Ouabain for 72 hours (by Presto Blue assay, n=3). Data were normalized to cell proliferation measured at 0 hours. (B) Percentage of PHA-stimulated $CD3^+$ T cells expressing CD69 and CD25 after 72 hours of co-culture with h-MSCs treated with Lubi (10 μM) or IVM (1 μM) (left panel) or AML-MSCs treated with 40 mM $K^+$ gluconate or 10 nM Ouabain (right panel), relative to untreated condition (n=3, p=not significant).

All histograms show mean ± SEM; *p-value <0.05, **p-value <0.01, ***p-value <0.001, ****p-value <0.0001.

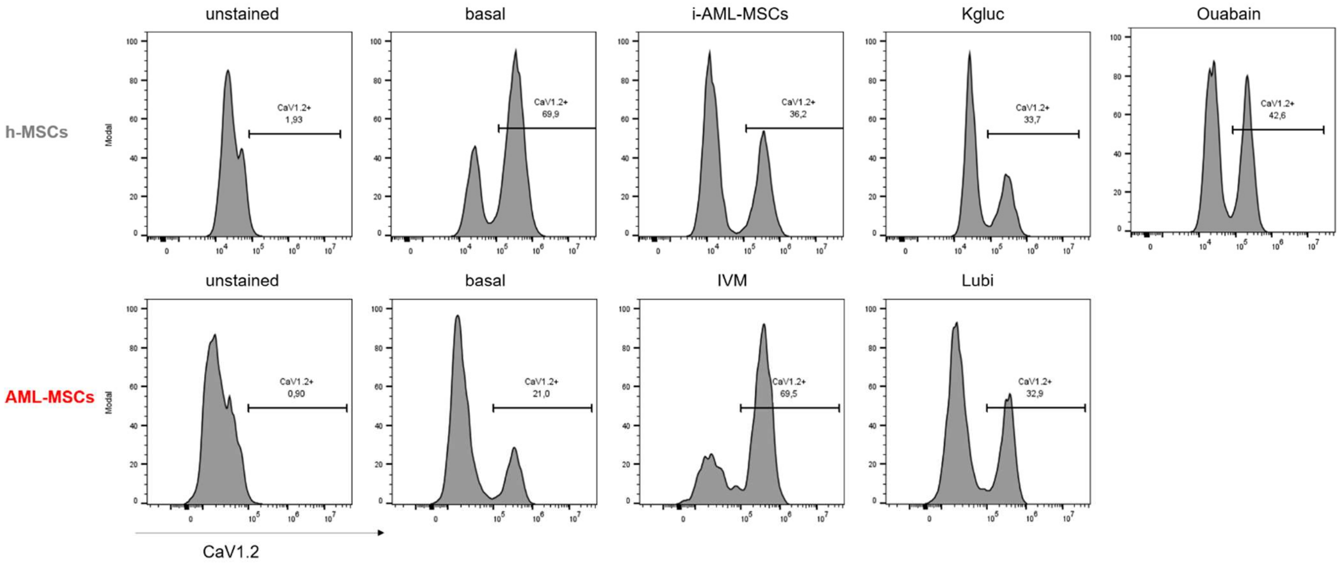


**Supplementary Fig. 5: $V_{mem}$ changes modify CaV1.2 expression levels.** Representative flow cytometry plots showing CaV1.2 expression on h-MSCs, h-MSCs co-cultured with AML blasts for 4 days (i-AML-MSCs) or h-MSCs treated with $K^+$ gluconate or Ouabain for 72 hours (upper panel). In the lower panel, histograms show CaV1.2 expression in AML-MSCs treated or not with Lubiprostone or IVM for 72 hours. Unstained samples are reported as control.

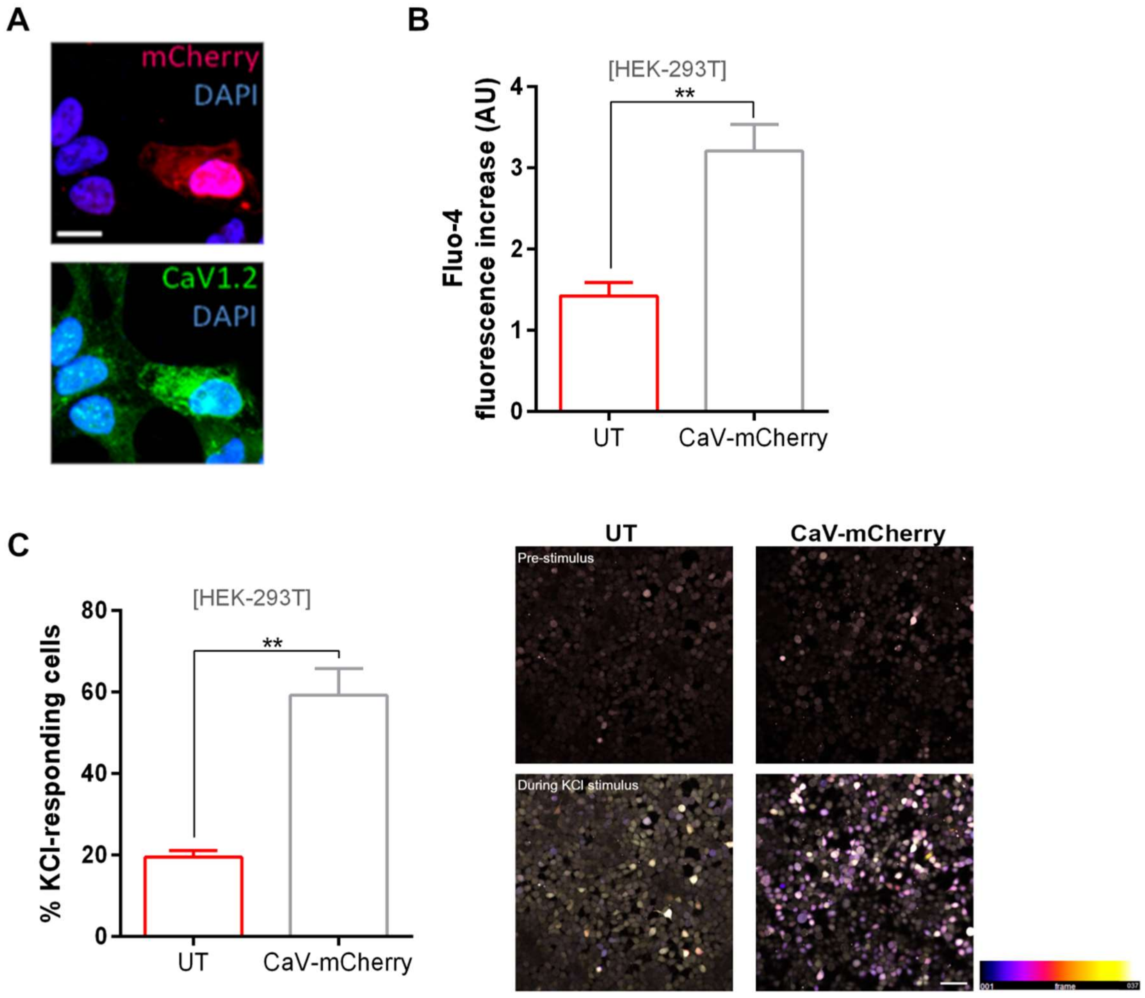


**Supplementary Fig. 6. Transient CaV1.2 overexpression controls calcium flux dynamics in HEK293T**. (A) Immunofluorescence staining on HEK293T cells with CaV1.2 antibody at confocal microscopy after 48 hours of transfection with CaV1.2-mCherry (red) plasmid. CaV1.2 and DAPI staining is green and blue, respectively (scale bar=10 µm, n=1). (B) Intracellular calcium increases in un-transfected HEK293T (UT, n=27 cells) and CaV1.2-mCherry-expressing cells (n=37 cells), loaded with Fluo-4 AM probe and stimulated with KCl (65 mM). C) Percentage of KCl-responding cells (left panel) and representative time sequence (pre- and during KCl stimulus) of intracellular calcium increase (right panel) in UT and CaV1.2-mCherry-expressing HEK293T cells loaded with Fluo-4 AM dye (n=4).

All histograms show mean ± SEM; ***p*-value <0.01.

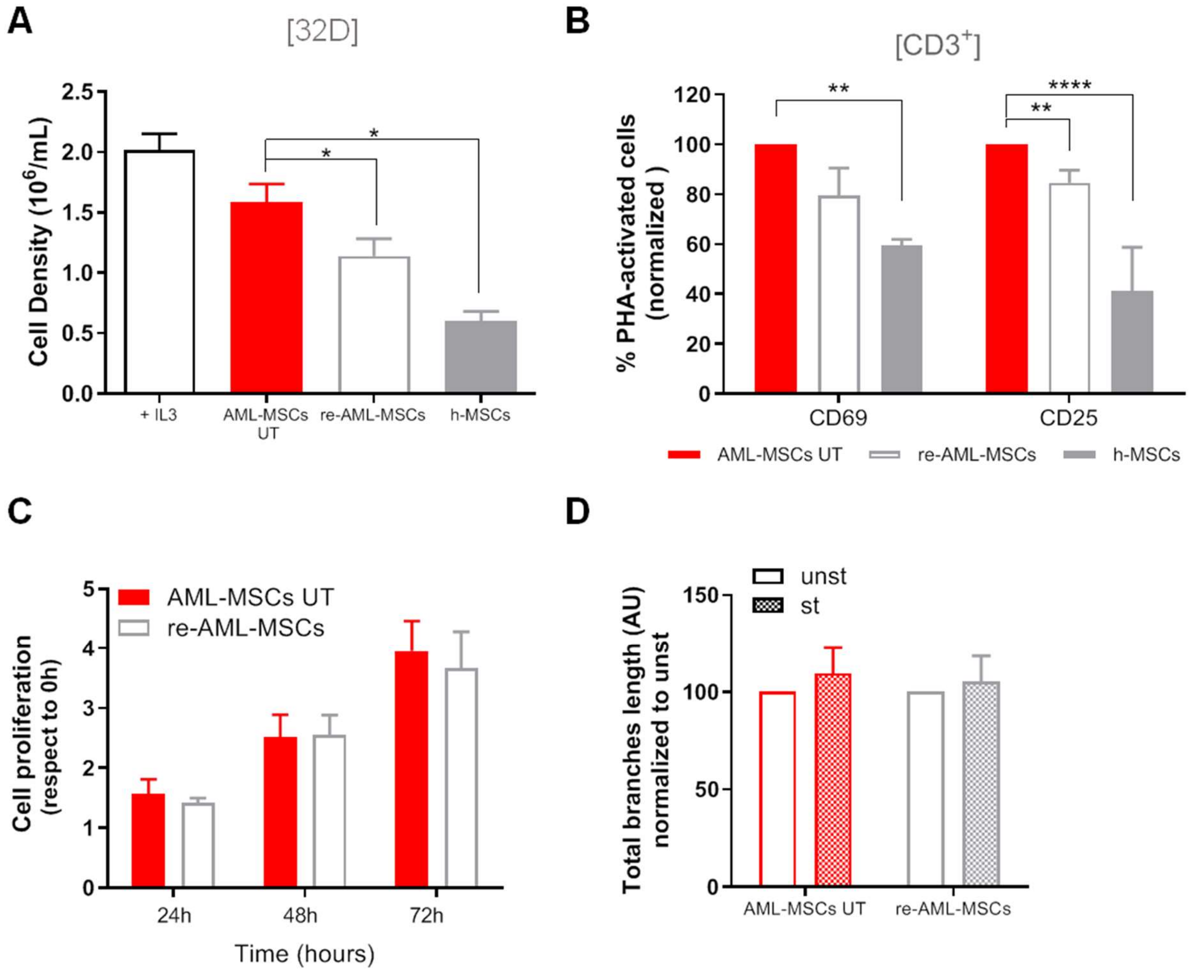


**Supplementary Fig. 7. CaV1.2 overexpression induces changes in MSCs phenotype.** (A) Cell density of murine IL-3–dependent 32D cell line cultured for 72 hours with IL-3 or on a layer of AML-MSCs UT, re-AML-MSCs (n=9) or h-MSCs (n=2, t-test comparing all groups *versus* AML-MSCs; +IL3 used as experimental control). (B) Percentage of PHA-activated T-cells expressing CD69 and CD25 markers after 72 hours of co-cultured with AML-MSCs UT, re-AML-MSCs (n=6) and h-MSCs (n=3). Data were normalized to AML-MSCs UT condition. (C) Cell proliferation measured in AML-MSCs UT and re-AML-MSCs for 72 hours by Presto Blue assay (n=7). (D) Total length of branches formed by HUVEC endothelial cells when cultured with conditioned medium derived from AML-MSCs UT or re-AML-MSCs (n=7) stimulated (st) or not (unst) with a pro-inflammatory cytokines cocktail for 24 hours; tube formation was evaluated after 4 hours and normalized to unst condition (AU: arbitrary unit).

All histograms show mean ± SEM; *p-value <0.05, **p-value <0.01, ****p-value <0.0001.

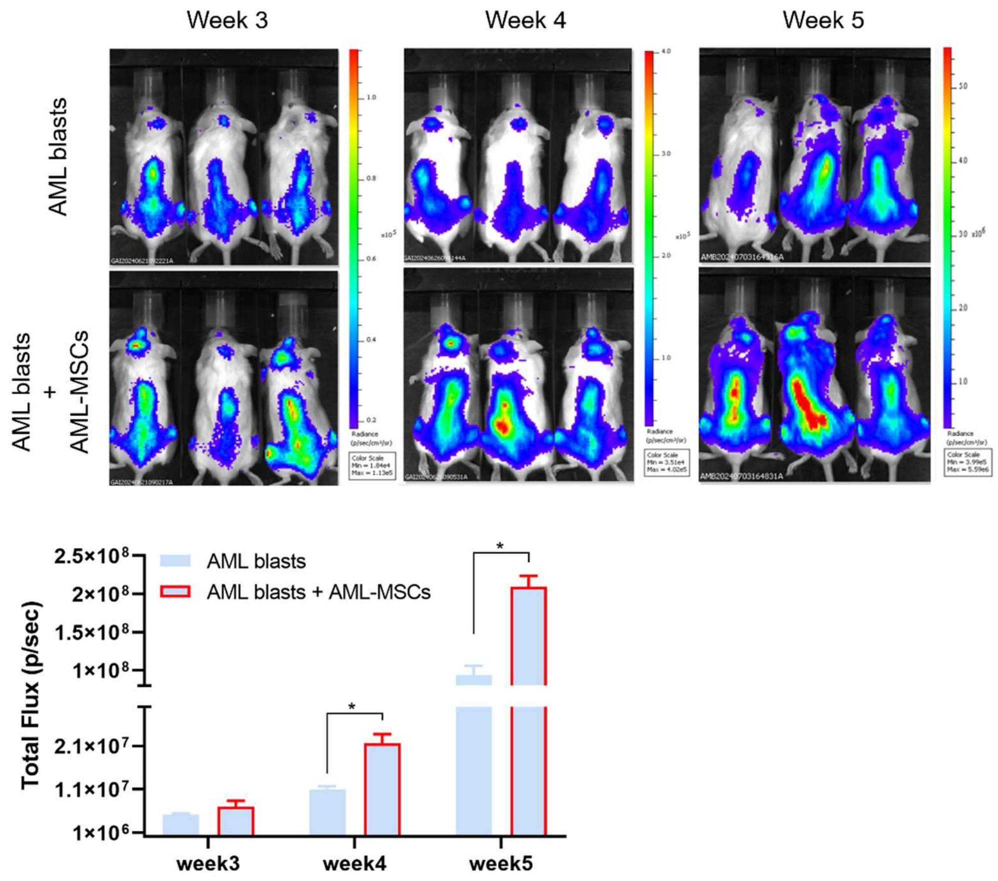


**Supplementary Fig. 8. Sustainment of AML-MSCs to blasts.** Tumor growth of luciferase-transduced primary AML cells injected in NSG mice with or without AML-MSCs, measured by luminescence activity (left panel). Images of bioluminescence in mice (right panel, n=3 mice per group). All histograms show mean ± SEM; *p-value <0.05.

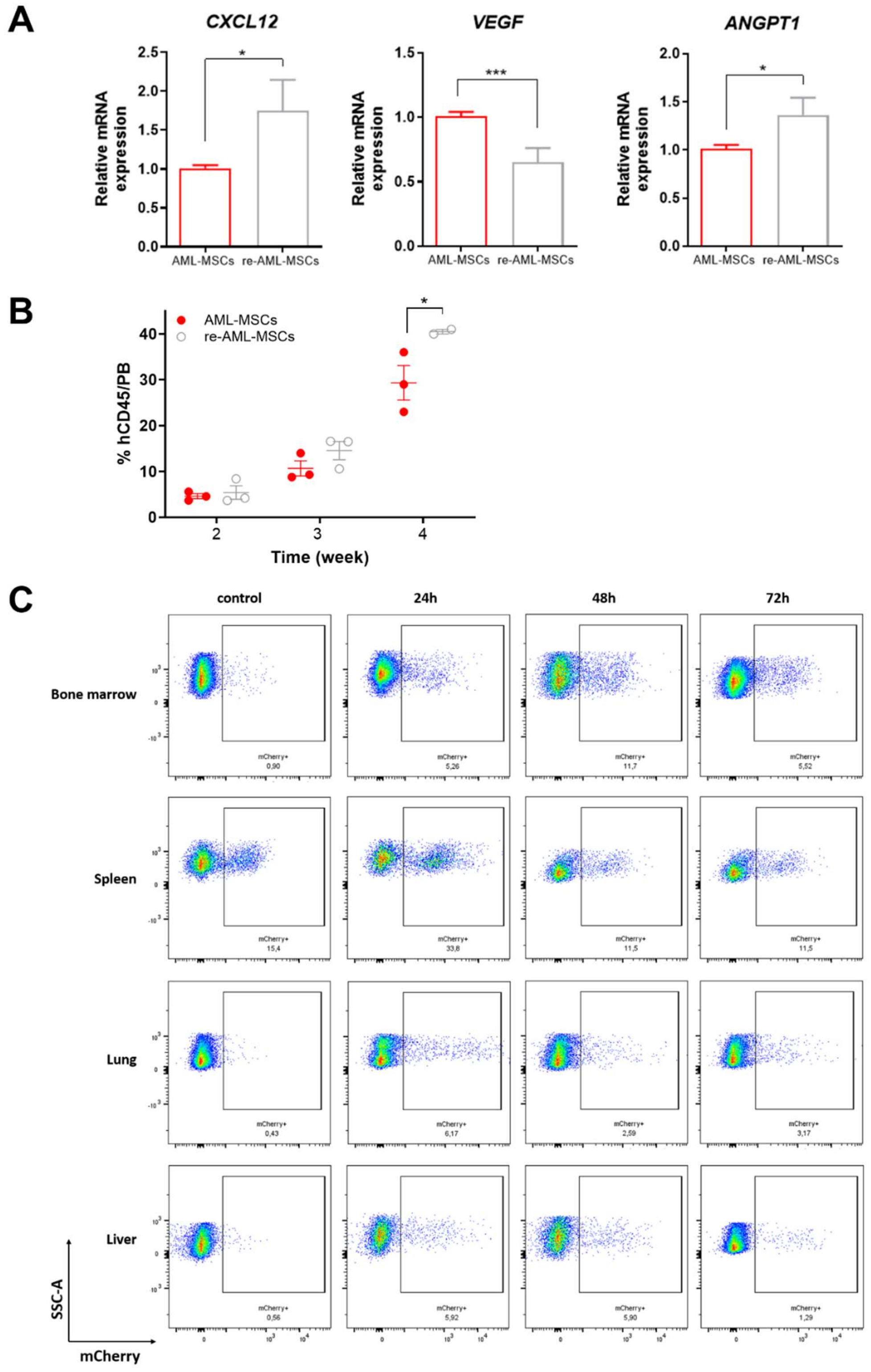


**Supplementary Fig. 9. MSCs influences normal hematopoiesis *in vivo*.** (A) Expression analysis of BM niche-associated genes (*CXCL12*, C-X-C motif chemokines ligand 12, n=6; *VEGFA*, vascular endothelial growth factor A, n=7; *ANGPT1*, angiopoietin 1, n=8) in AML-MSCs and re-AML-MSCs. Results are expressed as ΔΔCT relative to AML-MSCs UT control condition. (B) Percentage of human CD45 cells in NOG-EXL mice PB at 2, 3, and 4 weeks after tail vein coinfusion of $1.0x10^5$ human CB $CD34^+$ cells with $1.0x10^6$ AML-MSCs or re-AML-MSCs (n=3 mice per group). (C) Representative flow cytometry plots showing the tracking of mCherry-transduced MSCs in NSG mice in different districts (bone marrow, spleen, lung and liver) at 24-48-72 hours after injection.

All histograms show mean ± SEM; *p-value <0.05, ***p-value <0.001.

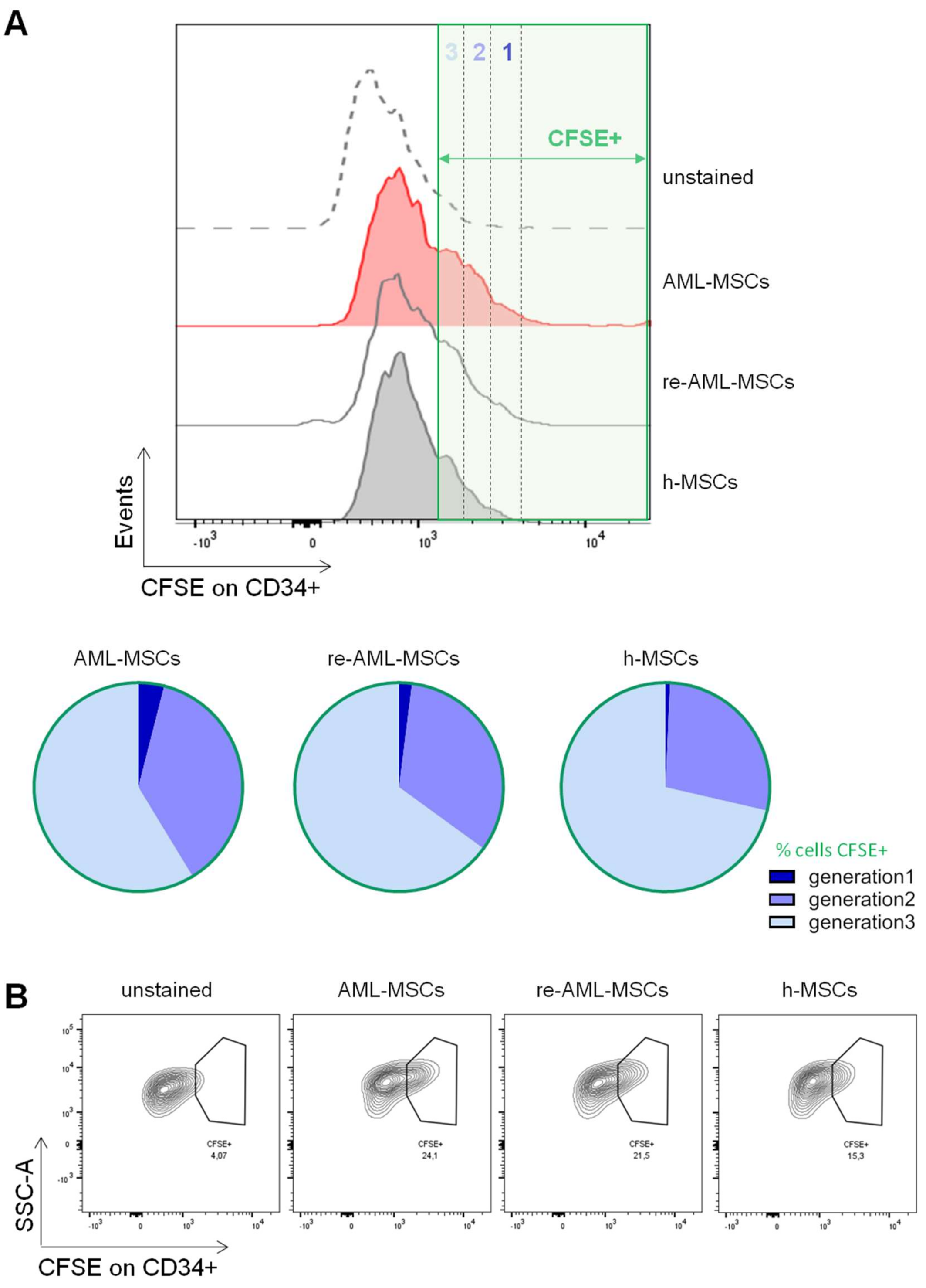


**Supplementary Fig. 10. Analysis of cell divisions in CD34+ cells by CFSE staining**. (A) CFSE staining of CD34+ cells harvested from bone marrow of mice co-injected with AML-MSCs, re-AML-MSCs or h-MSCs, 72 hours after injection, highlighting different proportion of cells in the different cell generations. (B) Analysis of CFSE+ CD34+ cells at 72 hours post-injection in mice with AML-MSCs, re-AML-MSCs or h-MSCs.
